# 3D-Printed Scaffolds Encapsulating Red Blood Cell Extracellular Vesicles for MicroRNA Delivery

**DOI:** 10.1101/2025.07.31.667777

**Authors:** Chongquan Huang, Migara Kavishka Jayasinghe, Vaibavi Ramanujam, Kieran Lau, Thi Nguyet Minh Le, Sing Yian Chew

## Abstract

Small non-coding RNAs (sncRNA) hold promising therapeutic potential. However, their clinical application is hindered by the poor cytocompatibility and limited transfection efficiency of conventional delivery vectors. In contrast, red blood cell-derived extracellular vesicles (RBCEVs) offer a safer, more efficient, and cost-effective alternative. Given the limited studies on the application of RBCEVs in the central nervous system (CNS) which is characterized by the presence of sensitive cell types with inherently low transfection efficiency, we hypothesized that RBCEVs could serve as a safe and effective sncRNA delivery vector for CNS applications, and that their incorporation into 3D-printed scaffolds could enable sustained and localized delivery of therapeutic sncRNAs. To test this, the uptake and gene silencing performance of RBCEVs were examined in primary CNS cell types, including astrocytes, neurons, oligodendrocyte precursor cells (OPCs), and microglia. While over 70% of OPCs and microglia internalized RBCEVs, uptake in neurons and astrocytes remained below 40%, indicating cell-type-specific uptake efficiency. Additionally, RBCEVs-mediated delivery of siRNA resulted in the highest gene knockdown efficiency in OPCs (74.2%), while triggering less than 30% gene knockdown in other cell types. Next, RBCEVs-encapsulated scaffolds were fabricated using digital light processing (DLP) 3D printing, enabling the sustained release of miR-219/miR-338-loaded RBCEVs for at least 21 days *in vitro*, which resulted in effective gene silencing that promoted OPC differentiation and myelination. Using spinal cord injury (SCI) as a proof-of-concept, scaffold-mediated delivery of RBCEVs-miR-219/miR-338 significantly promoted OPC differentiation and maturation *in vivo* as evidenced by increased CC1⁺ mature oligodendrocytes and reduced PDGFRα⁺ undifferentiated OPCs (*p* < 0.001). Taken together, these results demonstrate the therapeutic potential of combining RBCEVs with DLP-printed scaffolds for localized and sustained sncRNA delivery in CNS disease treatment.

## 1. Introduction

Small non-coding RNAs (sncRNAs), such as microRNAs (miRNAs) and small interfering RNAs (siRNAs), hold significant therapeutic potential by modulating gene expression with high specificity [1]. However, the clinical translation of sncRNA therapeutics remains significantly limited by the lack of safe and effective delivery systems [2].

The primary bottleneck in sncRNA delivery lies in the development of efficient and biocompatible vectors. Lipid nanoparticles and viral systems, although widely investigated in preclinical and clinical studies, are often associated with immunogenicity and cytotoxicity [3]. For instance, cationic liposomes and polymers have been reported to induce DNA damage, membrane destabilization, cell shrinkage, vacuolization, and activation of apoptotic pathways [4], while viral vectors frequently elicit severe immune responses [5]. In contrast, extracellular vesicles (EVs)-based delivery has emerged as a promising alternative due to their intrinsic biocompatibility and natural ability to transport cargos across cellular lipid bilayers [6]. Moreover, EVs possess surface ligands that facilitate active uptake and promote immune evasion, further enhancing their delivery efficiency compared to synthetic carriers [6]. Nevertheless, the clinical application of EVs faces challenges, including limited yields when derived from primary cells [7] and potential oncogenic risks associated with immortalized cell lines [8]. On the contrary, red blood cell-derived EVs (RBCEVs) offer a particularly attractive solution, benefiting from abundant blood availability, cost-effective production, and the absence of nuclear and mitochondrial DNA, thereby minimizing the risk of unintended nucleic acid-related side effects [9].

Beyond vector optimization, the administration route is equally critical to therapeutic success. Systemic delivery often leads to rapid clearance, off-target uptake, and associated side effects, necessitating high dosages to achieve therapeutic concentrations [10]. In contrast, scaffold-based delivery enables localized, sustained release while providing protection for the therapeutic agents against clearance and degradation [11]. Three-dimensional (3D) printing offers precise control over scaffold architecture and drug distribution, features that are essential for spatially confined and tunable release profiles. In particular, digital light processing (DLP)-based 3D printing achieves superior resolution, reproducibility, and production speed compared to conventional extrusion or laser-based methods, allowing accurate modulation of drug release kinetics [12]. With these advantages, DLP-printed scaffolds represent a promising yet underexplored platform for integrating RBCEVs to achieve localized, sustained, and spatially controlled delivery of therapeutic RNA cargos. Such a strategy is particularly relevant for treating complex pathological environments where conventional systemic delivery proves insufficient.

Current research has extensively explored the therapeutic potential of RBCEVs in cancer and related complications [13–16], while their application in the central nervous system remains largely underexplored, which requires a systematic investigation of uptake and transfection efficiency of RBCEVs in neural cells, such as astrocytes, neurons, oligodendrocyte precursor cells (OPCs) and microglia. Additionally, the use of scaffold-based delivery systems for RBCEVs remains underdeveloped, with limited studies addressing the long-term release dynamics, structural integration, and preservation of EVs bioactivity within 3D-printed platforms. These knowledge gaps largely restrict the development of stable, localized delivery systems for CNS disorders.

Spinal cord injury (SCI), characterized by acute inflammation, extensive secondary damage, and a hostile microenvironment, poses substantial challenges for therapeutic delivery, including rapid RNA degradation and drug clearance driven by dense infiltration of activated immune cells [17, 18]. Given these obstacles, SCI serves as an ideal proof-of-concept model to evaluate the delivery efficacy and therapeutic potential of this RBCEVs-integrated scaffold system. Here, we hypothesized that RBCEVs can effectively transfect CNS cells and be integrated into a 3D-printed scaffold via DLP 3D printing technology, while maintaining bioactivity, thereby exerting therapeutic effects.

## 2. Material and methods

### 2.1 Materials

DNase 1 type IV, Collagenase Type I (9001-12-1), L-cysteine, polycaprolactone (PCL), poly-d-lysine (PDL), Triton X-100, bovine serum albumin (BSA, A7030), citric acid, sodium citrate, glucose, dihydrogen phosphate, adenine, sodium chloride, mannitol, calcium ionophore, 4% trehalose Poly(ethylene glycol) diacrylate (PEGDA, average Mn 700), N-acetyl-l-cysteine (NAC),d-biotin, ITS, SATO components sodium dihydrogen phosphate, adenine and calcium chloride were purchased from Sigma-Aldrich. Neg miR (AM17110), miR-219 (PM10664), miR-338-3p (PM10716), and miR-338-5p (PM12825), Alexa-Fluor 633 goat anti-rabbit, Alexa-Fluor 633 goat anti-mouse, Alexa-Fluor 633 goat anti-rat antibodies, 4′,6-diamidino-2-phenylindole (DAPI), Dulbecco’s Modified Eagle Medium (DMEM) high glucose, neurobasal medium (NB), fetal bovine serum (FBS, 10,270,106), Glutamax (35,050,061), B-27 supplement (17,504,044), penicillin/streptomycin (Pen/Strep), trypsin-ethylenediaminetetraacetic acid (EDTA), Quant-iT RiboGreen kit, Pierce BCA Protein Assay, carboxyfluorescein diacetate succinimidyl ester (CFSE), and DPBS were purchased from Life Technologies. Rat anti-MBP (aa82-87) was purchased from Bio-Rad. Papain suspension was purchased from Worthington. Mouse anti-Integrin αM antibody (OX-42, CD11b) was purchased from Merck Millipore. 4% paraformaldehyde solution (PFA, sc-281,692) was purchased from Axil Scientific. Mouse anti-βIII Tubulin (Tuj1, 801,202) was purchased from Biolegend. Rabbit anti-GFAP (Z033401) was obtained from DAKO. Rabbit anti-Iba1 (019–19741) was purchased from Wako. Goat anti-PDGFRα (AF1062) was purchased from RND systems.Reversal transcript kit (NEB #E3010) and SYBR green qPCR kit (NEB #M3003) were purchased from New England Biolabs Inc. TransIT-TKO (MIR2150) was purchased from Mirus Bio, USA. The primers for Sox6, Hes5, PDGFRα, ZFP238, β-actin and GAPDH, the GAPDH siRNA (siGAPDH, IDT: rn. Ri.Gapdh.13.1) and negative siRNA (siNeg, 51-01-14-04) were predesigned and purchased from Integrated DNA Technologies Inc. (Coralville, IA). Leukoreduction filter was bought from Nigale, China. qEV-original size exclusion chromatography (SEC) column was purchased from Izon, New Zealand. RNeasy Mini Kit was purchased from Qiagen. Gelatin methacryloyl (GelMA) and Lithium phenyl-2,4,6-trimethylbenzoylphosphinate (LAP) were purchased from EFL, China. EVfect transfection kit was purchased from Jotbody HK Ltd.

### 2.2 RBCEVs Preparation and Characterization

Human RBCEVs were isolated from fresh human red blood cells (RBCs) in collaboration with ESCO Aster (Singapore), following protocols outlined in our prior work[16]. Human red blood cells were obtained from donors with informed consent, provided by Innovative Research Inc. (USA). Briefly, RBCs were separated from whole blood through 3 rounds of centrifugation and washing in PBS (1000g for 8 min at 4 °C) to eliminate platelets and plasma. The enriched RBCs were then resuspended in PBS and filtered through a leukoreduction filter to remove any residual leukocytes. The resulting RBC suspension was then collected in Nigale buffer, consisting of 0.2 g/L citric acid, 1.5 g/L sodium citrate, 7.93 g/L glucose, 0.94 g/L sodium dihydrogen phosphate, 0.14 g/L adenine, 4.97 g/L sodium chloride, and 14.57 g/L mannitol, and subsequently diluted in CPBS (PBS with 0.1 mg/mL calcium chloride). To promote vesiculation, the RBCs were incubated overnight with calcium ionophore (10 μM final concentration) at 37 °C and 5% CO₂. Following vesiculation, human RBCEVs were isolated by sequential differential centrifugation to remove cell debris, filtered through a 0.45 μm filter, and then concentrated by ultracentrifugation at 21,000g for 30 min. A sucrose cushion ultracentrifugation step was employed to reduce protein contamination, and the resulting EV pellet was washed twice with PBS. After purification, the RBCEVs were resuspended in PBS supplemented with 4% trehalose and stored at −80 °C. All experiments conducted in this study utilized human RBCEVs. The purity of RBCEVs was verified using western blot against specific EV markers, namely Band3 and Stomatin. The relative abundance of EV markers was assessed between RBCEVs and RBCs by the blot as well.

To track RBCEVs uptake and biodistribution, RBCEVs were labelled with CFSE which yield stable fluorescent signals in the presence of esterases (which are also present within the lumen of RBCEVs). CFSE was dissolved in anhydrous DMSO to prepare a 5 mM stock solution. RBCEVs were prepared in PBS at a final concentration of 1 mg/mL, and the CFSE stock was added to achieve a working concentration of 20 μM. The RBCEVs were incubated with the CFSE dye at 37 °C for 60 min to ensure complete activation and fixation of the dye inside the RBCEVs, followed by centrifugation at 21,000g to pellet the labeled RBCEVs. The RBCEVs pellet was resuspended and subsequently washed twice at 21,000g for 30 min to remove any free unbound CFSE dye. The labelled RBCEVs suspension was subsequently passed through a qEV-original SEC column to further purify the labelled RBCEVs and ensure the complete removal of any free CFSE dye. Fractions 7-9 from the SEC column containing RBCEVs were collected and concentrated by centrifugation at 21,000g at 4 °C for 20 min. These purified CFSE-labelled RBCEVs were used for subsequent downstream experiments.

Purified RBCEVs were loaded with nucleic acid payloads using the EVfect transfection kit following the manufacturer’s instructions. Loaded and unloaded RBCEVs were characterized using nanoparticle tracking analysis with a ZetaView nanoparticle tracking instrument (Particle Metrix). The loading efficiency of nucleic acid into RBCEVs was semi-quantified via electrophoresis by comparing nucleic acid content on loaded RBCEVs to a serial dilution of nucleic acid.

### 2.3 Primary CNS Neural Cell Isolation

The isolation of primary rat CNS neural cells was carried out following our published protocol [19] and in accordance with the guidelines of the Institutional Animal Care and Use Committee (IACUC, Protocol number: A240001) at Nanyang Technological University. Briefly, P0–P2 neonatal rat cortices were isolated and enzymatically digested with 1.2 U papain, 0.24 mg/mL L-cysteine and 40 μg/mL DNase at 37 °C for 1 h. Thereafter, the digestion was stopped by adding DMEM full media (DMEM with 10% FBS and 1% Pen/Strep). The dissociated tissues were centrifuged at 300g for 5 min. Thereafter, the tissue pellets were used for downstream processing.

#### 2.3.1 Primary Cortical Neuron Isolation and Culture

After adding NB full media (Neurobasal media with 10% FBS, 2% B27 supplement, 1% GlutaMAX, and 1% Pen/Strep) to the cell pellet from Section 2.3, the pellets were gently pipetted 15 times to form a homogeneous single cell suspension. Thereafter, the cell suspension was filtered through a 70 μm cell strainer and centrifuged at 300g for 5 min. The cell pellets were then resuspended and seeded at a density of 1.5 x 10^5^ cells/well in a 24-well plate pre-coated with 20μg/mL PDL. Following that, cells were maintained with NB full media without FBS, at 37 °C, 5% CO2.

#### 2.3.2 Primary Mixed Glial Culture and Glial Cell Separation

The cell pellets from Section 2.3 were resuspended with DMEM full media and triturated 10 times with a 21 G needle and another 10 times with a 23 G needle. After obtaining a single-cell suspension, cells from 6 digested cortices were seeded onto four PDL-coated (2 μg/mL) T75 flasks and cultured in DMEM full media with media change every 2–3 days until Day 10.

<u>Microglia culture:</u> At Day 10, flasks were shaken on an orbital shaker at 200 rpm for 1 h at 37 °C. The supernatants containing loosely attached microglia were centrifuged at 300g for 5 min, and the microglial cell pellet was collected. The cell pellets were resuspended and then seeded at a density of 2 x 10^5^ cells/well in a PDL-coated 24-well plate or PDL pre-coated 13-mm round coverslip. Thereafter, the cells were maintained with DMEM full media, at 37 °C, 7.5% CO_2_.

<u>OPCs culture:</u> After removal of microglia, the flasks were refilled with DMEM full media for another 16 h of shaking at 200 rpm, 37 °C. The supernatants consisted largely of OPCs, with small amounts of microglia. The remaining microglia were removed by incubating the supernatants on untreated petri dishes for 25 min, allowing the microglia to attach due to their rapid adhesion properties. Thereafter, the supernatants were centrifuged at 300g for 5 min and the OPCs pellet was used for downstream experiments.

Proliferation media (DMEM with 100x Sato, 1% Pen/Strep, 1% ITS supplement, 0.5% FBS,10 ng/ml PDGF and 10 ng/ml FGF2/bFGF) was used for short-term cultures of OPCs for cellular uptake and gene silencing assays as depicted in Sections 2.5 and 2.6, respectively. Specifically, cells were seeded at a density of 1.5 x 10^5^ cells/well in a PDL-coated 24-well plate or PDL pre-coated 13-mm round coverslip. To evaluate the extent of oligodendrocyte myelination *in vitro*, myelination media (DMEM/Neurobasal = 50:50, with 2% B27, 1% Pen/Strep, 0.5% Glutamax, 1% ITS, 10 ng/mL biotin, 5 μg/mL NAC, and 1% SATO) was used for myelination culture with cells seeded at a density of 20,000 cells/ cm^2^ on scaffolds that comprised of suspended electrospun PCL fibers that were fabricated according to our published protocol [20]. Throughout all experiments, cells were maintained at 37 °C, 7.5% CO_2_.

<u>Astrocyte culture:</u> After the removal of OPCs, astrocytes that remained on the flasks were collected by treatment with 0.25% trypsin for 20 min at 37 °C. The digestion was then halted by adding DMEM full media at a volume ratio of 1:1. After centrifugation at 300g for 5 min, cell pellets were collected, resuspended and seeded with DMEM full media at a density of 7.5 x 10^4^ cells/well in a PDL-coated 24-well plate or PDL pre-coated 13-mm round coverslip. Cells were then maintained at 37 °C, 7.5% CO_2_.

### 2.4 *In Vitro* Cytocompatibility Assays

<u>Live-dead assay:</u> Cells with 500 μL of media were incubated with 25 μg of RBCEVs for 24 h, washed with DPBS, and replenished with fresh media. 20 μM of Calcein-AM and Propidium iodide, diluted 1:1000 (v/v) in DPBS, (1 μL Calcein-AM + 1 μL Propidium iodide + 998 μL DPBS) were added to the cell media and incubated for 30 min at room temperature. Following 2x PBS washing, cells were refilled with media and imaged immediately. More than 3 biological replicates were conducted.

<u>CCK-8 metabolic assay:</u> After removal of media, cells were washed with DPBS twice. Fresh media with 10% CCK-8 (30 μL CCK-8 in 270 μL media) was added into each well and incubated for 30 min at 37 °C. Thereafter, the supernatant was collected for measurement using a microplate reader at 450 nm absorbance. More than 3 biological replicates were conducted.

### 2.5 *In Vitro* RBCEV Cellular Uptake Assay

Individual CNS cell types were cultured in 500 μL of media, at the density as mentioned in section 2.3.1 and 2.3.2 in 24-well plates. After 24 h, either 25 or 50 μg of CFSE-RBCEVs, were added into cell culture wells and incubated for another 24 h before processing.

<u>Lysotacker assay and live-cell imaging:</u> Cells were incubated with 50 nM lysotracker-488 diluted with corresponding culture media (as mentioned in section 2.3.1 and 2.3.2) for 1 h. Thereafter, cells were washed with PBS 3 times and the labeled Cy5-ODN RBCEVs were added into cell culture refilled with fresh media and imaged using a Carl Zeiss Cell Discoverer 7 for 24 h. Lysoendosomal escape was then calculated by:

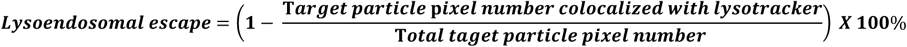

<u>Flow cytometry:</u> Cells were washed with PBS before treatment with 200 μL/well of 0.25% trypsin. After 20 min incubation, 200 μL of DMEM full media was added to deactivate trypsin. The media-trypsin mixture was collected and centrifuged at 300g for 5 min, followed by removal of the supernatant. Cells were then resuspended with pre-chilled 0.5% BSA in DPBS and taken for flow cytometry analysis. FITC channel was used to detect the cells that internalized CFSE-RBCEVs. Three biological replicates were conducted.

<u>Cell immunostaining:</u> Cells were washed with PBS before fixation with 4% PFA. After washing off the PFA, permeabilization and blocking were performed by incubating all samples in blocking buffer, which comprised of 1% BSA and 0.1% Triton in PBS for 1 h. Specific CNS cell marker antibodies (Microglia: Iba1, 1:500; Astrocyte: GFAP, 1:1000; OPCs: PDGFRα, 1:500; Neuron: Tuj1, 1:1000) diluted with 1% BSA in 0.1% PBST (0.1% Triton in PBS) were added to each well respectively, and incubated at 4 °C overnight for primary staining. The following day, samples were washed 3 times with PBS. Thereafter, relevant Alexa Fluor-tagged secondary antibodies (1:1000) and DAPI (1:1000) were diluted with PBS and added correspondingly to each sample. All samples were incubated for 1 h. Following that, cells cultured on 13-mm diameter round coverslips were fixed on top of a rectangular coverslip with mounting media for imaging under an epifluorescence microscope (Leica DMi8) or a confocal microscope (Zeiss, LSM800). More than 3 biological replicates were conducted.

### 2.6 Gene Transfection

For bolus transfection with siGAPDH, each CNS cell type was cultured in 24-well plates at cell densities as described above. For each well, 0.5 μL of siGAPDH (100 μM) or siNeg (100μM) was mixed with 0.5 μL of TransIT-TKO reagent and another 9 μL of nuclease-free water and incubated at room temperature for 10 min. Thereafter, the mixture was added to one well with 490 μL of media. For transfection using siGAPDH-RBCEVs or siNeg-RBCEVs, 25 μg of RBCEVs (500 ng RNA) was added into each well with 500 μL media. Thereafter, cells were maintained for 48 h before RNA extraction.

Similarly, for miR-219/miR-338 transfection, 0.167 μL of miR-219, 0.167 μL of miR-338-3p, and 0.167 μL of miR-338-5P stock (100μM) were mixed with 0.5 μL of TKO and another 9 μL of nuclease-free water and incubated at room temperature for 10 min. 0.5 μL of Neg miR was mixed with 0.5 μL of TKO and another 9 μL of nuclease-free water and incubated at room temperature for 10 min. Thereafter, the mixture was added to one well with 490 μL of media. For transfection using miR-219/miR-338-RBCEVs or Neg miR-RBCEVs, 25 μg of RBCEVs (500 ng RNA) was added into each well with 500 μL of media. Following that, cells were maintained for 48 h before taken for qPCR processing and analysis.

The scaffold-based transfection was done using a trans-well culture system (800 μL media/ well), but the process varied depending on the purpose. OPCs cultured on PDL-coated well plate (as described in Section 2.3.2) were co-incubated alongside the scaffolds placed in the trans-well inserts for 48 h before RNA extraction for gene expression analysis. OPCs fiber suspension cultures (also described in Section 2.3.2) were also co-incubated alongside scaffolds that were placed in the trans-well inserts and cultured for 3, 10 or 14 days, with media changes every 2 days. At each timepoint, cells were fixed and taken for immunostaining to analyze the extent of oligodendrocyte differentiation and myelination.

### 2.7 qPCR

After 48 h of transfection, total RNA was isolated and purified using the RNeasy Mini Kit (Qiagen) based on the manufacturer’s protocol. Thereafter, reverse transcription was conducted following the manufacturer’s instructions. Specifically, 200 ng of total RNA was added with 4 μL of RE mix and water to obtain a final total volume to 20 μL. The synthesizing process was set as 30 min to obtain cDNA. For qPCR analysis, SYBR green master mix was used with a StepOnePlus™ real-time PCR system (Applied Biosystem), with β-actin serving as the housekeeping gene. Each reaction mix included 1 μL of cDNA and 0.25 μL of 5 μM primers as shown in Supplementary Table 1. The qPCR was carried out using a two-step program consisting of 15 s of denaturation followed by 30 s of extension, repeated for 40 cycles. The comparative delta Ct method was used to analyze gene expression levels, after ensuring that all primers had similar amplification efficiencies. Each condition was tested in at least 3 biological replicates with 3 technical replicates.

### 2.8 Scaffold Fabrication

RBCEVs-incorporated scaffolds were fabricated using DLP 3D printing. Specifically, the ink comprised of 10% w/v GelMA, 5% w/w PEGDA, 500 μg/mL RBCEVs and 0.2% w/v LAP dissolved in PBS. The ink was crosslinked using a single printing layer at 25 mW/cm^2^ for 8 s to form the scaffold using an Asiga MAX X27. After printing, each scaffold was incubated with 200 μL of PBS (pH = 7.4) at 37 °C for 1 h to remove any remaining un-crosslinked ink.

### 2.9 Release Kinetics Study

After washing as mentioned at Section 2.8, the scaffolds (∼5 mg, with outer dimensions of 3 mm in diameter and 2 mm in height) were incubated in 200μL PBS at pH 7.4 and 37 °C for 1, 2, 3, 5, 7, 10, 14, and 21 days. At each timepoint, all the supernatant was collected for analysis and refreshed with 200 μL fresh PBS. RBCEVs particle number was measured using a DLS nanosizer (Malvern Zetasizer Nano). To quantify the miR content, the supernatant was treated with 0.1% Triton-X for 15 min to release all miRs from RBCEVs. The amount of miRs that was released into the supernatant was then quantified using the Quant-iT RiboGreen assay, following the manufacturer’s protocol. The total loading amount was determined by summing the quantity of RBCEVs or RNA released into the supernatant as described above, and the amount retained within the scaffold, which was measured after dissolving the residual scaffold using 100 U/mL collagenase at 37°C for 20 minutes.

### 2.10 Animal Surgery and Tissue Processing

The Institutional Animal Care and Use Committee (IACUC) of Nanyang Technological University (NTU) approved all animal procedures under protocol number A20015.

All animal experiments were conducted on female Sprague-Dawley rats (7 - 9 weeks, 250 - 280 g). Anesthesia was induced by intraperitoneal injection of ketamine (75 mg/kg) and xylazine (10 mg/kg). The surgical area was prepared by following the sequence of shaving, washing with 70% ethanol, and wiping with betadine. Thereafter, the T8-T11 vertebrae was exposed by surgery. A dorsal laminectomy was performed at the T9-T10 level. Following that, 2 mm of the spinal cord was removed. The 3D printed scaffold was then placed in the resulting gap. The muscles were sutured, and the skin was closed using wound clips. At 2 weeks and 4 weeks post-transplantation, the animals underwent perfusion with 0.9% saline to remove blood. Thereafter, 4% PFA was used for perfusion. The spinal cord with scaffold was isolated and further fixed with 4% PFA for more than 1 day before being transferred to 15% sucrose for an additional 24 h. Subsequently, the samples were soaked in 30% sucrose overnight for cryoprotection. Spinal cord samples were sectioned longitudinally at a thickness of 20 µm using a cryostat. Thereafter, the samples were directly mounted onto superfrost glass slides (Leica). Immunostaining for each cell type was conducted for the tissue samples to evaluate cellular responses. Specifically, cryosection tissue slides were permeabilized with 0.3% Triton in PBS for 10 min, followed by washing with PBS for 3 times, before blocking with 1% BSA in 0.1% PBST for 1 h. Primary antibodies, including CC1 (1:500), PDGFRα (1:800), iNOS (1:200), CD206 (1:500), OX42 (1:800), and Iba1 (1:1000), diluted with 1% BSA in 0.1% PBST were then added and the samples were incubated at 4 °C overnight. Following that, all samples were washed with PBS thrice before incubation with secondary antibodies corresponding to each primary antibody species (all at 1:800 diluted with PBS) for 1 h at room temperature. After washing with PBS for 3 times, samples were stained with DAPI (1:1000, diluted with PBS) for 5 min followed by PBS washing for another 3 times. Finally, samples were dried and mounted with mounting media for imaging.

### 2.11 Statistical Analysis

Normality and homogeneity of variances were assessed using Levene’s test. One-way ANOVA followed by Tukey’s post-hoc test was applied to data with normal distribution and equal variances. For non-normally distributed data or those with unequal variances, Kruskal-Wallis and Mann-Whitney U tests were used for comparisons involving more than two groups. Student’s t-test was performed for comparisons between two independent samples. Data were reported as mean ± standard deviation (SD).

## 3. Results

### 3.1 RBCEVs are cytocompatible with primary neural cells from the CNS

RBCEVs were successfully isolated from RBC as characterized in Figure S1A. RBCEVs-specific markers, Stomatin and Band3, were detected in RBCEVs samples but absent in RBCs as tested by western blot, showing successful isolation. Figure S1B shows that RNA loading did not change the size distribution, which peaked at ∼200 nm as measured by DLS nanosizer.

To evaluate the potential of RBCEVs for sncRNA delivery to neural cells, their cytocompatibility was compared to PBS (blank control) and TransIT-TKO, a commercial transfection reagent used in our previous studies [21–24]. As shown in Figure 1, live/dead staining after 24 h of treatment showed that RBCEVs did not induce significant cytotoxicity in primary astrocytes, neurons, OPCs and microglia, with live cell ratios comparable to those observed in PBS-treated cells. In contrast, a significantly higher level of OPC death was induced by TransIT-TKO vs. PBS- and RBCEVs-treatment. Additionally, RBCEVs treatment did not induce metabolic activities changes in neurons and glial cells (Figure S2). Collectively, these results demonstrate that RBCEVs are cytocompatible with primary neural cells and exhibit superior cytocompatibility as compared to TransIT-TKO, particularly in OPCs.

**Figure 1.**
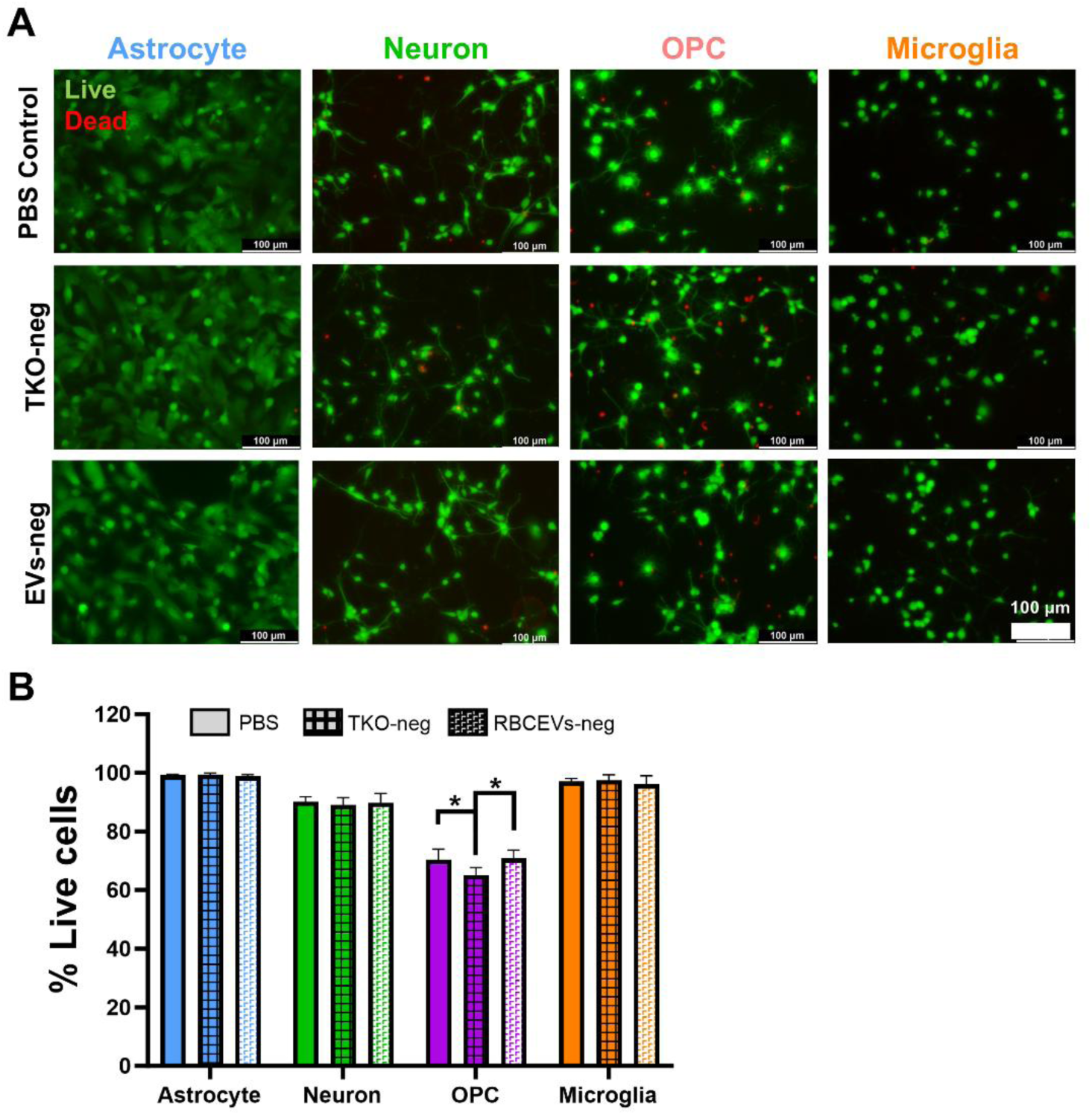
Cytocompatibility of RBCEVs with primary CNS-derived neural cells. (A) RBCEVs did not induce cytotoxicity, as shown by live/dead staining of astrocytes, cortical neurons, OPCs, and microglia treated with 25 μg of EVs for 24 h; (B) Quantitative analyses suggest that RBCEVs did not trigger significant cell death in cortical neurons and glial cells, although TransIT-TKO triggered increased death of OPCs. \**p* < 0.05, Brown-Forsythe test with one-way ANOVA, n = 3 cell batches

### 3.2 RNA-loaded RBCEVs successfully transfected primary CNS-derived neural cells and induced silencing of target genes

As shown in Figure 2A, confocal imaging confirmed the successful internalization of CFSE-labeled RBCEVs by primary neurons and glial cells after 24 h of treatment. Furthermore, as indicated in Figure 2B, a cell type-specific uptake preference of RBCEVs was observed, with the highest internalization seen in microglia and OPCs (90.1% and 76.0%, respectively), vs. neurons (37.0%) and astrocytes (23.2%).

**Figure 2.**
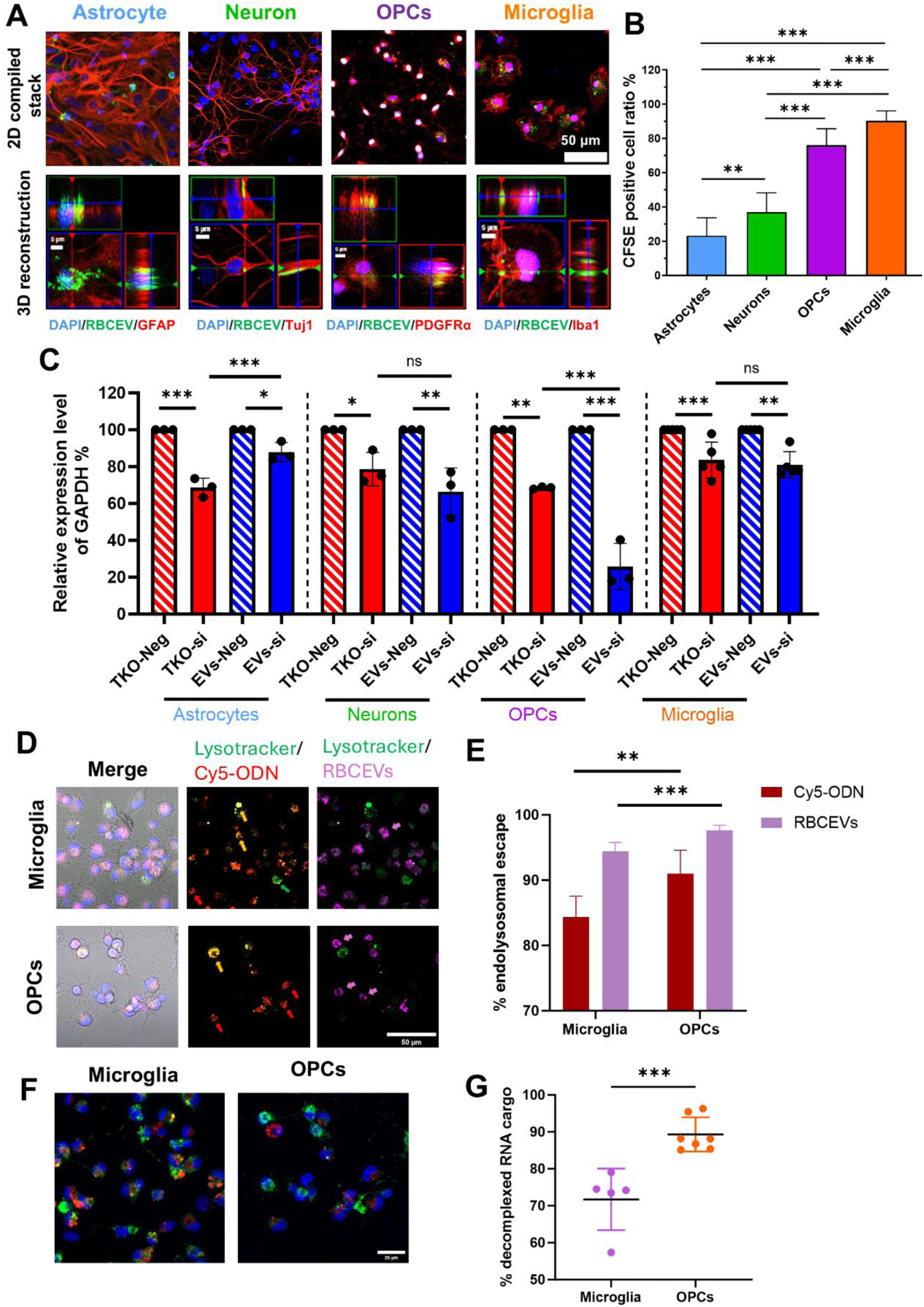
RBCEVs successfully entered CNS cells and triggered gene knockdown with cell type-specific efficiency. (A) Cytoplasmic localization of RBCEVs in primary CNS cells. Blue: DAPI; Green: CFSE-labeled RBCEVs; Red: cell-specific markers (GFAP: astrocytes, Tuj1: neurons, PDGFRα: OPCs, Iba1: microglia). Top: confocal z-stack maximum projection. Bottom: magnified orthogonal view; (B) Quantification of RBCEV uptake ratios showed significantly higher uptake in microglia, followed by OPCs, neurons, and astrocytes; (C) qPCR analysis revealed that 48 h treatment of TKO-siGAPDH or RBCEV-delivered siGAPDH produced comparable silencing in microglia and neurons, whereas TKO was more effective in astrocytes and RBCEVs were more effective in OPCs; (D&E) Colocalization pattern of Cy5-ODN to lysotracker revealed a higher extent of endolysosomal escape of ODN and RBCEVs in OPCs than in microglia. Red arrow: non-colocalized Cy5-ODN; yellow arrow: colocalized Cy5-ODN; magenta arrow: non-colocalized RBCEVs; green arrow: non-colocalized endolysosomes; (F&G) OPCs exhibited a greater fraction of decomplexed Cy5-ODN (red pixels) out of the total Cy5-ODN signal (yellow plus red pixels) than microglia. Statistical analysis: Brown-Forsythe test with one-way ANOVA (B) and unpaired t-test (C,E&G); \*\*\**p* < 0.001, \*\**p* < 0.01, \**p* < 0.05; n = 3 cell batches.

The RNA transfection capacity of RBCEVs in neural cells was also evaluated by qPCR, with TransIT-TKO as the positive control. As shown in Figure 2C, TransIT-TKO triggered significantly more extensive gene knockdown in astrocytes than RBCEVs (*p* < 0.001), while RBCEVs significantly induced > 2-fold gene knockdown in OPCs than TransIT-TKO (*p* < 0.001). In microglia and neurons, the knockdown efficiencies between RBCEVs and TransIT-TKO were comparable. Taken together, these results suggest that the knockdown efficiency varied depending on the delivery vehicle and cell type, and highlighted OPCs as an ideal target for RBCEVs-based RNA delivery.

To understand the potential mechanisms associated with the observed higher uptake but lower gene knockdown efficiencies in microglia than OPCs, the extent of endolysosomal escape was assessed in these 2 types of cells. As shown in Figure 2D, compared to microglia, OPCs showed larger proportion of free Cy5-ODN and RBCEVs at 48 h post-transfection, as indicated by a higher number of red or magenta particles not overlapping with green endolysosomal markers. Furthermore, quantitative analysis of the extent of endolysosomal escape revealed that OPCs exhibited a greater extent of escape for Cy5-ODN (∼91%) and RBCEVs (∼98%) than microglia (∼84% vs. ∼94%, Figure 2E). These results indicate that RBCEVs and ODN exhibited greater endolysosomal escape in OPCs than in microglia, resulting in increased cytoplasmic functional siRNA and thereby enhanced gene knockdown.

### 3.3 miR-219/miR-338-loaded RBCEVs successfully induced targeted gene silencing in OPCs

Given their superior RNA transfection efficiency, OPCs were selected to assess the therapeutic potential of RBCEVs. To examine dose-dependent uptake, OPCs were treated with CFSE-labeled RBCEVs at 25 μg/well or 50 μg/well. Correspondingly, fluorescence imaging and FACS analysis showed increased signal intensity and CFSE⁺ cell numbers at the higher dose, with FACS revealing an increase from 53.0% to 69.4% (Figure 3A).

**Figure 3.**
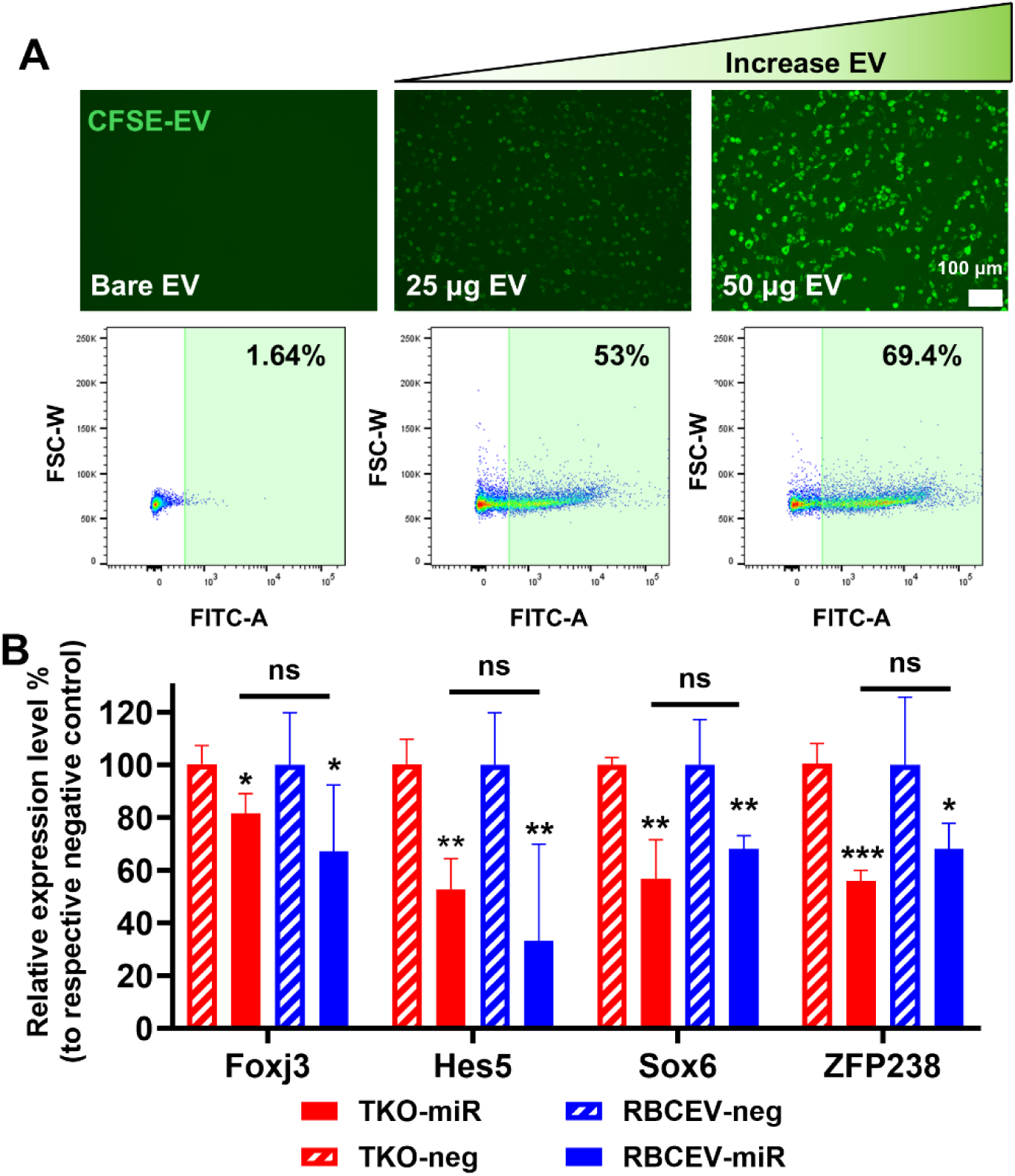
Successful uptake and miR-mediated gene silencing in OPCs by RBCEVs. (A) RBCEV uptake in OPCs increased with higher RBCEV concentrations, as shown by elevated CFSE-positive cell ratios and fluorescence intensities in imaging and flow cytometry; (B) RBCEVs loaded with miR-219/miR-338 triggered comparable knockdown of Foxj3, Hes5, Sox6, and ZFP as TransIT-TKO despite lower dosage (500 ng in RBCEVs vs. 600 ng in TransIT-TKO). Statistical analysis was conducted between TKO-miR/TKO-neg as well as between RBCEV-miR/RBCEV-neg by unpaired t-test, \*\*\**p* < 0.001, \*\**p* < 0.01, \**p* < 0.05; n = 3 cell batches.

MiR-219 and miR-338 regulate oligodendrocyte differentiation and myelination through the repression of targets, including Foxj3, Hes5, Sox6, and Zfp238 [25]. Building on these findings, our previous study applied the TransIT-TKO system to deliver miR-219/miR-338 for therapeutic promotion of post-injury remyelination [21]. To compare the efficacy of RBCEVs over TransIT-TKO in delivering miR-219/miR-338 to OPCs, the target downstream genes were analyzed by qPCR at 48h post-transfection. As shown in Figure 3B, treatment with miR-219/miR-338-loaded RBCEVs effectively decreased the expressions of Foxj3, Hes5, Sox6, and Zfp238. Despite containing less microRNA in RBCEVs (500 ng versus 633 ng in TransIT-TKO), RBCEVs achieved comparable gene knockdown to that of TransIT-TKO (*p* > 0.05), indicating a higher transfection efficiency. Altogether, these findings support the potential of RBCEVs as an effective delivery system for miRs, particularly targeting OPCs.

### 3.4 DLP printing enabled incorporation and sustained release of bioactive miRs-loaded RBCEVs from 3D-printed scaffolds

RBCEVs were encapsulated into microchannel scaffolds by DLP printing. As demonstrated in Figure 4A, successful incorporation of RBCEVs was achieved with uniform drug distribution throughout the scaffolds. As shown in Figure 4B, RBCEVs released from scaffolds incubated at 37 °C and pH 7.4 for 2-48 h maintained a size distribution profile comparable to freshly prepared non-encapsulated RBCEVs. These results indicated that the structural integrity of RBCEVs was well preserved after both the printing process and their subsequent sustained release from the scaffolds.

**Figure 4.**
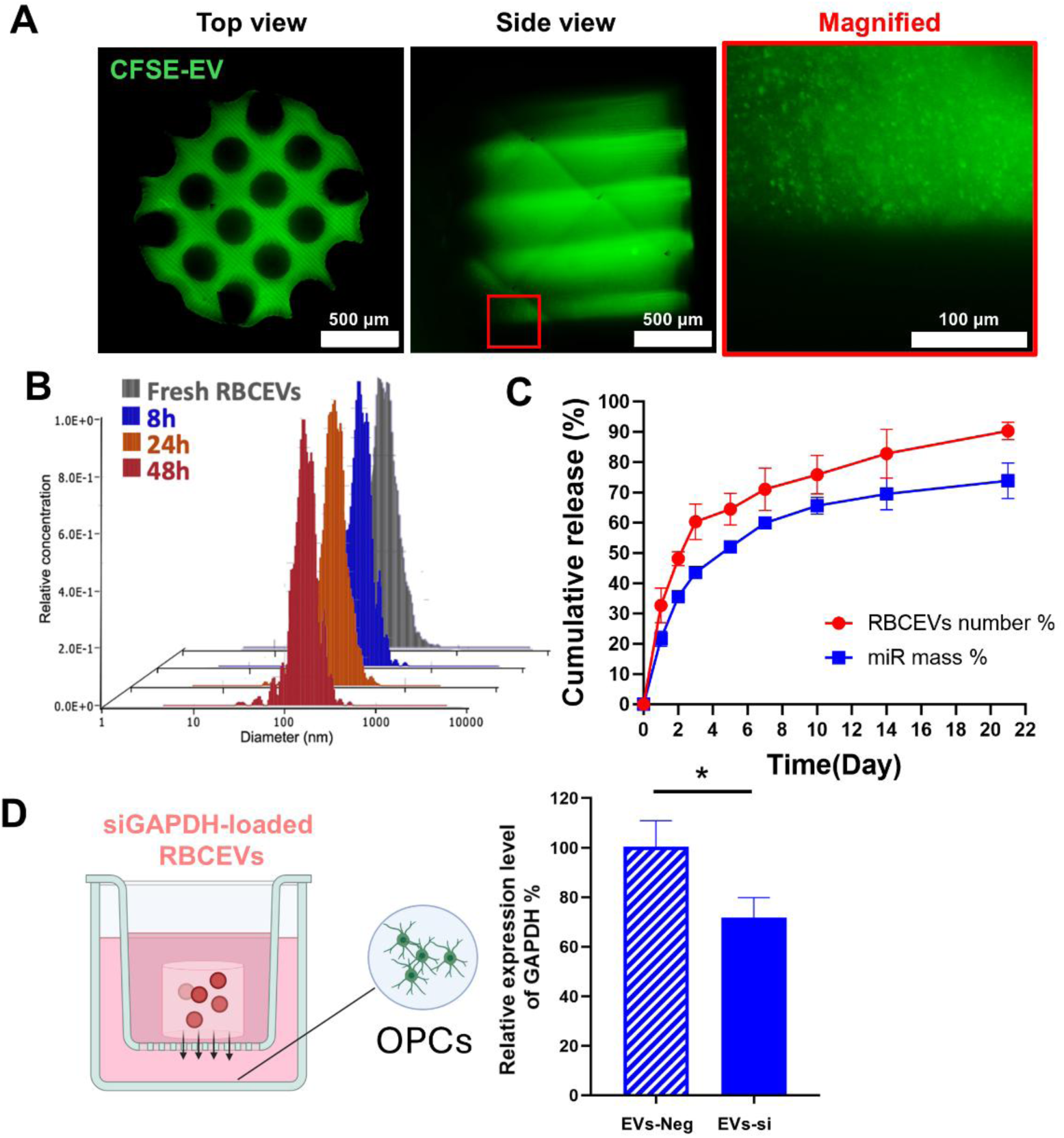
Encapsulated RBCEVs retained structural and functional integrity after 3D printing and showed sustained released from 3D-printed scaffolds. (A) Top, side, and magnified views of a 3D-printed scaffold encapsulated with CFSE-RBCEVs, demonstrating uniform RBCEV distribution; (B) Size distribution of fresh vs. released RBCEVs suggested structural stability for at least 48 h *in vitro*; (C) Release kinetics analysis showed that both RBCEVs and miRs were sustainably released from 3D-printed scaffolds for at least 21 days in vitro, based on either percentage of theoretical loading. n = 3 scaffolds; (D) qPCR analysis of OPCs co-cultured with scaffolds encapsulating siNEG- or siGAPDH-loaded RBCEVs demonstrated that siGAPDH delivery significantly reduced GAPDH expression. Statistical analysis: unpaired t-test; \**p* < 0.05; n = 3 cell batches.

As shown in Figure 4C and Figure S3, a sustained release of RBCEVs and miRs was obtained for at least 21 days *in vitro*. In particular, after an initial burst release of (32.6 ± 5.8) % and (21.6 ± 2.5) % of RBCEVs and miRs respectively (i.e. 246.9 ± 5.2 ng RBCEVs/mg of scaffold and 2.3 ± 0.1 ng miRs/mg of scaffold), (90.3 ± 2.9) % and (73.8 ± 5.9) % of RBCEVs and miRs (i.e. 858.2 ± 172.3 ng RBCEVs/mg of scaffold and 23.0 ± 5.5 ng miRs/mg of scaffold) were released respectively by Day 21. These results demonstrate a consistent ratio between RBCEVs and miRs, suggesting the well-preserved integrity of RBCEVs.

The gene silencing capacity of the RBCEVs-encapsulated scaffolds was further evaluated. As shown in Figure 4D, scaffolds incorporated with siGAPDH-loaded RBCEVs induced significant gene knockdown of GAPDH in OPCs as compared to siNeg-RBCEVs-incorporated scaffolds (*p* < 0.05), indicating that the RBCEVs remained functional after encapsulation and release from the scaffolds, effectively triggering gene knockdown.

### 3.5 Scaffolds incorporating miR-219/miR-338-loaded RBCEVs enhanced OPC differentiation and myelination *in vitro*

To examine whether RBCEVs-encapsulated scaffolds could effectively deliver miRs and regulate OPCs activities, miR-219/miR-338- or scrambled, non-functional Neg miR-loaded RBCEVs were incorporated into scaffolds and co-incubated with OPCs. OPCs were cultured on a suspending-fiber platform as elaborated in Figure 5A.

**Figure 5.**
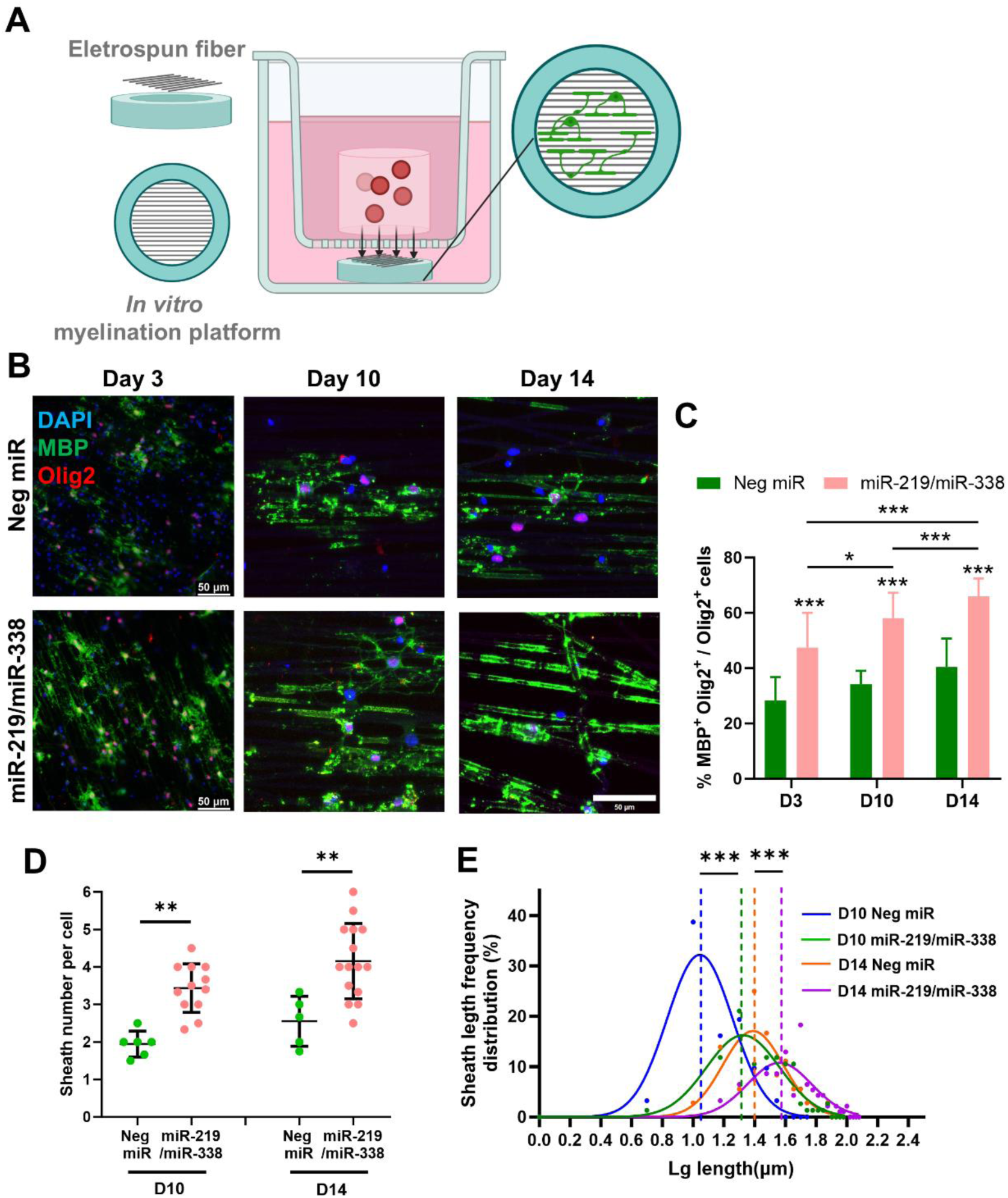
Scaffolds encapsulating miR-219/miR-338-loaded RBCEVs enhanced OPC differentiation and myelination in vitro. (A) Scheme of OPCs myelination culture platform and co-incubation with scaffold. (B) Fluorescence images showing OPCs seeded on aligned PCL fibers in the presence of scaffolds, which contained RBCEVs loaded with either Neg miR or miR-219/miR-338 in a transwell system at Days 3, 10, and 14. Blue: DAPI; Green: MBP; Red: Olig2; (C) miR-219/miR-338 induced higher percentage of MBP+ mature oligodendrocytes at D3, D 10 and D 14; (D) Sheath number per cell and (E) sheath length demonstrated improved oligodendrocyte myelination with miR-219/miR-338-RBCEVs. Statistical analysis: Brown-Forsythe test with one-way ANOVA; \*\*\**p* < 0.001, \*\**p* < 0.01, \**p* < 0.05; n = 3 cell batches.

As shown in Figure 5B and 5C, miR-219/miR-338 treatment significantly increased the number of MBP⁺ mature oligodendrocytes at Day 3, 10, and 14 as compared to Neg miR-treated OPCs. While a modest rise in MBP⁺ cells was observed over time with Neg miR treatment, it did not reach statistical significance. In contrast, miR-219/miR-338 induced a robust, time-dependent enhancement in OPCs differentiation and maturation. These findings demonstrate the efficient delivery and functional activity of miR-219/miR-338 from the DLP-printed scaffolds in promoting the differentiation and maturation of OPCs.

Importantly, miR-219/miR-338 treatment also promoted oligodendrocyte myelination at Day 10 and Day 14. Specifically, miR-219/miR-338 treatment resulted in a significantly higher number of myelin sheaths formed per cell compared to negative control miRs (Figure 5D). Frequency distribution analysis of sheath lengths revealed a significant shift toward longer myelin sheaths at both Day 10 and Day 14 following miR-219/miR-338 treatment (Figure 5E). Moreover, a significant time-dependent increase in sheath length was observed from Day 10 to Day 14 no matter treated with neg miR or miR-219/miR-338, indicating the spontaneous myelination properties of OPCs culture on fibrous substrate.

### 3.6 Scaffold-mediated delivery enabled efficient and spatially localized delivery of RNA-loaded RBCEVs *in vivo*

Localized and sustained drug delivery could enhance therapeutic efficacy while minimizing systemic side effects. As shown in Figure 6A, 1 week after implantation of the scaffolds, CFSE-labeled RBCEVs were distributed up to 3 mm caudally and 2.5 mm rostrally from the host–implant interface, whereas Cy5-ODN remained primarily within 1 mm from the interface, indicating localized delivery. Additionally, CFSE signals in the scaffold area remained detectable after 7 days of implantation, suggesting sustained release of RBCEVs from the scaffolds.

**Figure 6.**
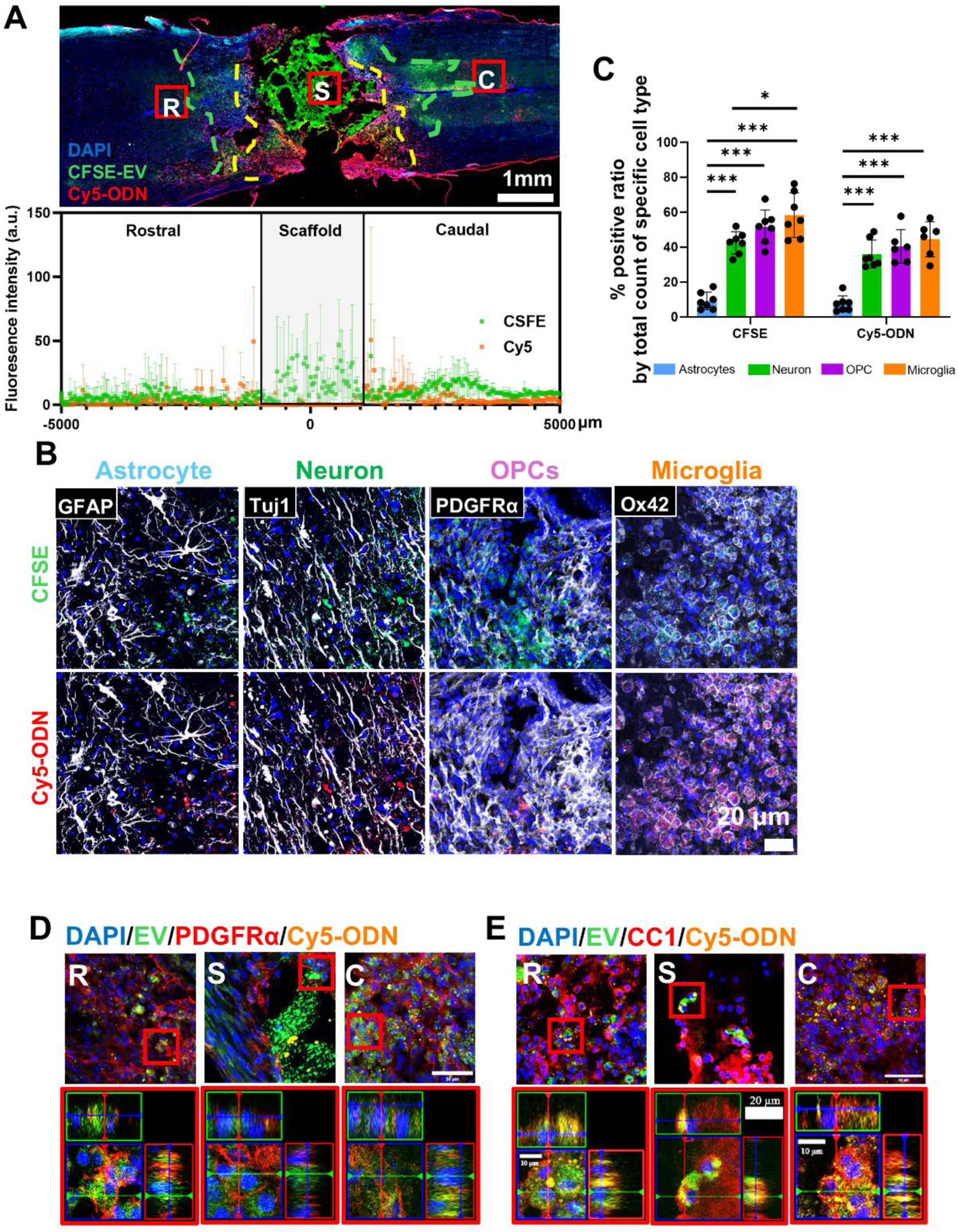
Scaffolds encapsulating RBCEVs achieved highly efficient, spatially localized delivery to OPCs and oligodendrocytes *in vivo*. (A) Representative fluorescent image of spinal cord tissue implanted with a scaffold showed RBCEVs (green) and Cy5-ODN (yellow) uptake, peaking within ∼2.5 cm rostrally and 3 cm caudally. The curve graph quantifies fluorescence intensity along the spinal cord. R, S, and C represents rostral, scaffold and caudal region of spinal cord section, respectively. The dotted line represents the furthest extent of CFSE-RBCEVs and Cy5-ODN distribution; (B) Representative images demonstrating in vivo delivery of CFSE-RBCEVs and Cy5-ODN from scaffolds to OPCs, microglia, neurons and astrocytes. ROIs were selected within 100 μm from tissue-scaffold interface; (C) RBCEVs and ODN predominantly were delivered to neurons, OPCs, and microglia, while their delivery to astrocytes was minimal. Statistical analysis: Brown-Forsythe test with one-way ANOVA; \**p* < 0.05, \*\*\**p* < 0.001, n = 3 animals x 5 ROI.; (D)&(E) Representative magnified images of the rostral, scaffold, and caudal regions of the spinal cord showing successful uptake of RBCEVs and Cy5-ODN in PDGFR+ OPCs (D) and CC1+ mature oligodendrocytes (E). Orthogonal views (bottom, red frames) suggested intracellular localization of RBCEVs and Cy5-ODN. Blue: DAPI; Green: CFSE-RBCEVs; Red: PDGFRα (D) or CC1 (E); Yellow: Cy5-ODN. Original image size: 160 × 160 μm, scale bar = 50 μm; magnified image size: 30 × 30 μm.

The ability of the scaffolds in delivering sncRNAs into cells *in vivo* was subsequently evaluated. As shown in Figures 6B and 6C, CFSE-labeled RBCEVs and Cy5-labeled RNA were observed in neurons and glial cells marked by cell-type-specific markers, confirming effective cellular internalization. Specifically, the uptake of RBCEVs and delivery efficiency of RNA displayed a cell-type-specific pattern, with highest efficiency in microglia and OPCs, followed by neurons, and astrocytes (Figure 6C). In particular, a significantly lower uptake was seen in astrocytes relative to the other cell types (*p* < 0.001).

Based on the RBCEVs distribution pattern shown in Figure 6A, further analysis was performed in 3 regions: one inside the scaffold, and the other two at the distal rostral and caudal extreme of RBCEVs distribution. As shown in Figures 6D and 6E, both Cy5-ODN and RBCEVs were detected within the cytoplasm of PDGFRα⁺ OPCs and CC1⁺ mature oligodendrocytes, indicating that the scaffold enabled effective delivery of sncRNA to oligodendroglial cells across different developmental stages.

### 3.7 Scaffold-mediated delivery of miR-219/miR-338-loaded RBCEVs promoted OPCs differentiation and maturation *in vivo*

To assess the effect of miR-219/miR-338 loaded RBCEVs-encapsulated scaffolds on OPCs fate regulation *in vivo*, key oligodendroglial lineage markers were analyzed. Figure 7A shows a marked reduction in PDGFRα expression following treatment with miR-219/miR-338, indicating a decrease in undifferentiated OPCs compared to the Neg miR treated group. This observation was supported by quantification in Figure 7B, which reveals a more than 75% reduction in the PDGFRα⁺ Olig2⁺/Olig2⁺ cell ratio under the influence of miR-219/miR-338 vs. Neg miR treatment (*p* < 0.001).

**Figure 7.**
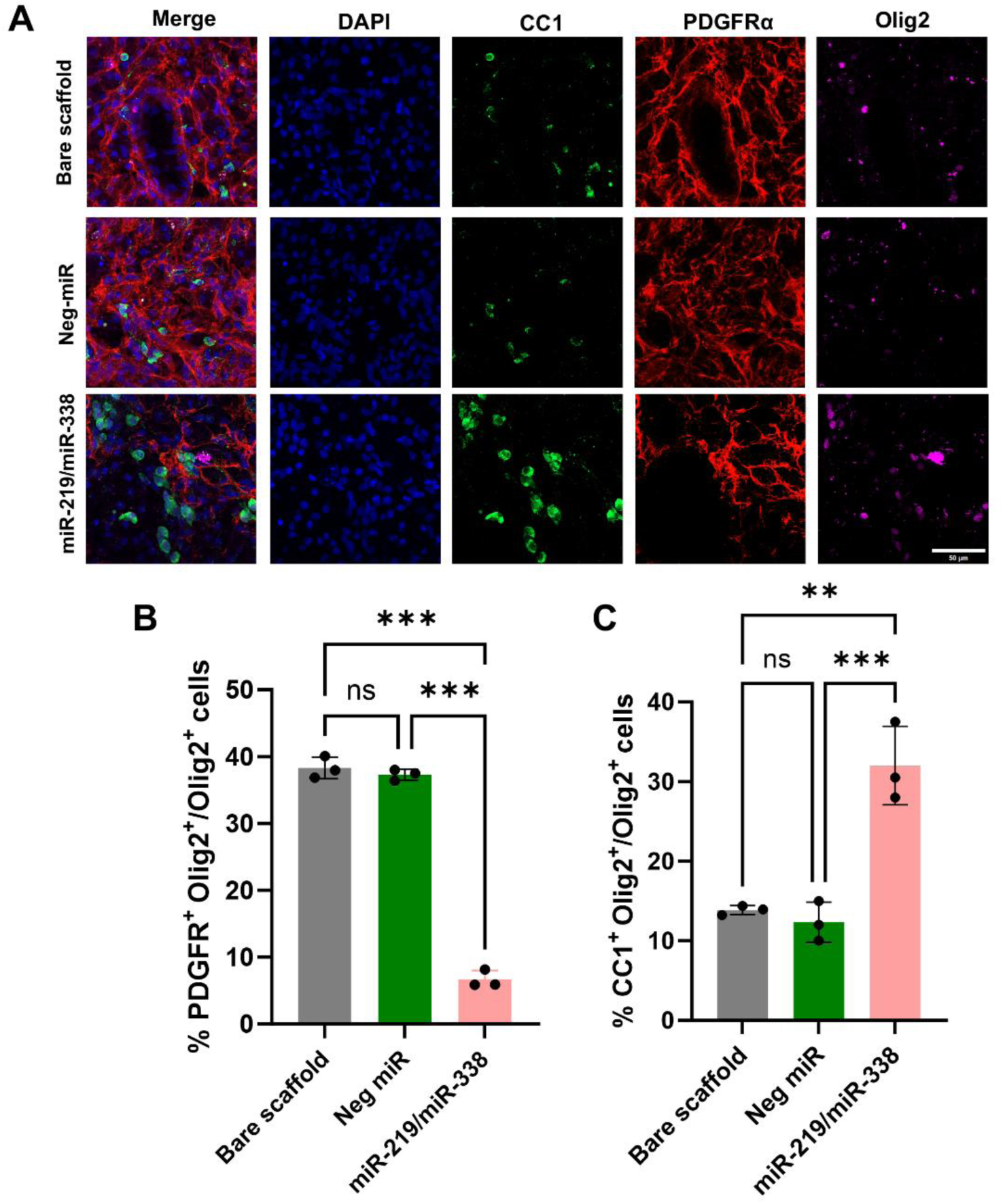
Scaffolds encapsulated with miR-219/miR-338-loaded RBCEVs promoted OPCs differentiation and maturation. (A) Representative images showing that miR-219/miR-338 promoted OPC differentiation and maturation in vivo, as evidenced by fewer PDGFRα⁺ OPCs and more CC1⁺ mature oligodendrocytes when compared to bare scaffold or Neg-miR scaffold. ROIs were selected within 100 μm from the tissue-scaffold interface. Scale bar = 50 μm; (B & C) Quantitative analysis of PDGFRα⁺Olig2⁺/Olig2⁺ and CC1⁺Olig2⁺/Olig2⁺ ratios showing enhanced OPCs maturation with miR-219/miR-338 RBCEVs scaffold treatment. One-way ANOVA; \*\**p* < 0.01, \*\*\**p* < 0.001; n = 3 animals.

In contrast, expression of CC1, a marker of mature oligodendrocytes, was notably enhanced in the miR-219/miR-338-treated group. As shown in Figure 7A and quantified in Figure 7C, the CC1⁺ Olig2⁺/Olig2⁺ cell ratio increased by over 2-fold when animals were treated with miR-219/miR-338, as compared to Neg miR treatment (*p* < 0.001), indicating an expansion of the mature oligodendrocyte population.

Altogether, these results demonstrate that miR-219/miR-338-loaded RBCEVs scaffolds effectively promoted the *in vivo* transition of OPCs towards a more differentiated oligodendroglial state.

### 3.8 Scaffold encapsulated with miR-219/miR-338-loaded RBCEVs did not cause detrimental side effects

The immune response was assessed by examining microglial polarization markers. Confocal imaging (Figure 8A) showed that scaffolds loaded with miR-219/miR-338 RBCEVs significantly reduced the proportion of iNOS⁺ pro-inflammatory microglia as compared to both the bare scaffolds (without RBCEVs) and the Neg miR-loaded scaffolds, as confirmed by quantification in Figure 8B.

**Figure 8.**
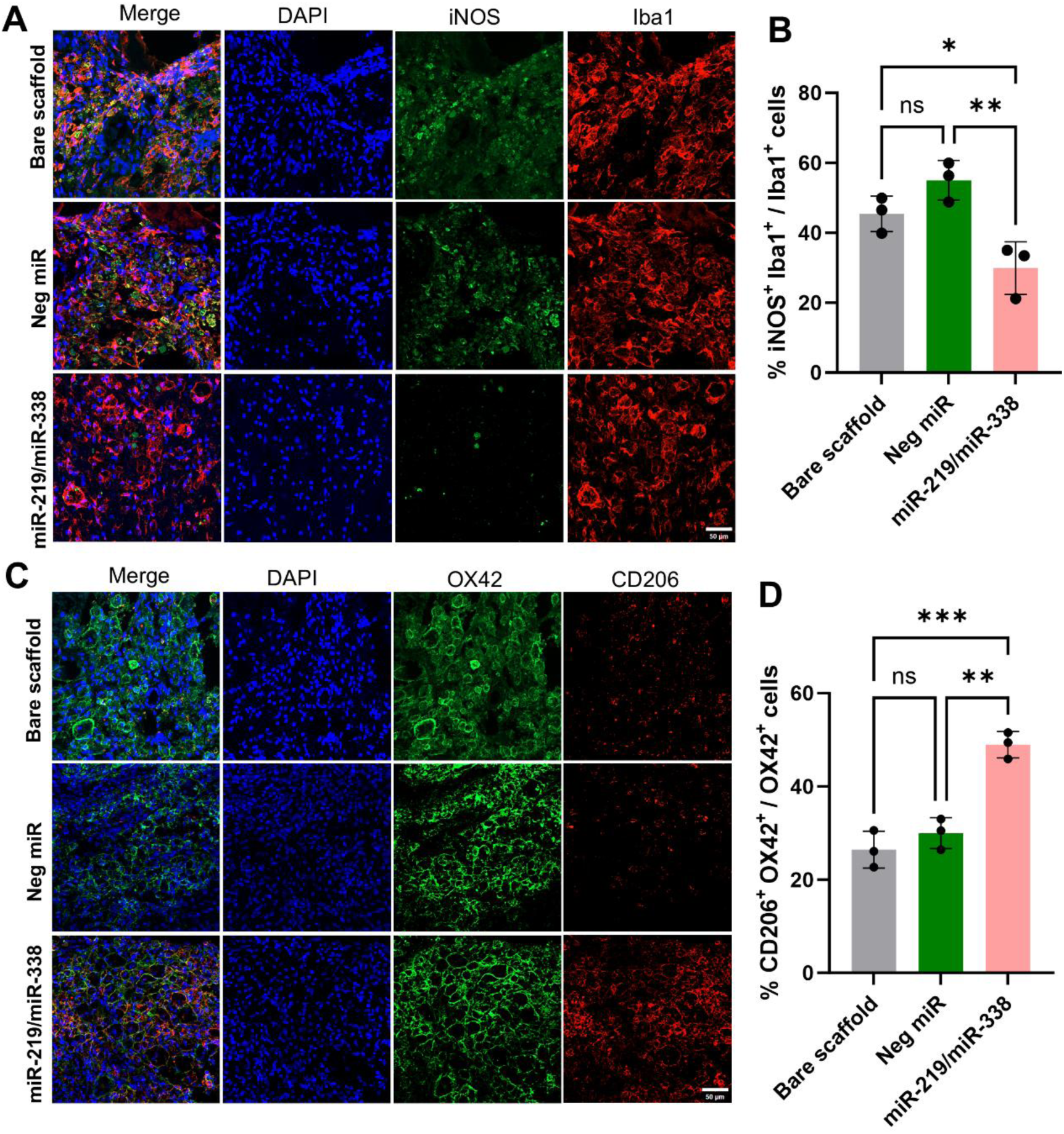
Scaffolds encapsulated with miR-219/miR-338-loaded RBCEVs induced anti-inflammatory phenotype of microglia. (A&B) Representative fluorescence images and quantification showing that scaffolds loaded with miR-219/miR-338 RBCEVs significantly reduced the proportion of pro-inflammatory (iNOS⁺) microglia among total Iba1⁺ cells. No significant difference was observed between the bare scaffolds and the Neg miRNA-loaded scaffolds. (C&D) Representative fluorescence images and quantification showing that miR-219/miR-338 RBCEV-loaded scaffolds significantly increased the proportion of anti-inflammatory (CD206⁺) microglia among total OX42⁺ cells. No significant difference was observed between the bare scaffolds and the Neg miRNA-loaded scaffolds. ROIs were selected within 100 μm from tissue-scaffold interface. Scale bar = 20 μm; Statistical analysis: One-way ANOVA; ns: no significance; *p* > 0.05, \**p* < 0.05, \*\*\**p* < 0.001; n = 3 animals.

In addition, a significantly higher proportion of CD206⁺ anti-inflammatory microglia was observed in the miR-219/miR-338 group relative to both bare scaffold and Neg miR scaffolds (*p* < 0.01), indicating the anti-inflammatory effect of the miRs.

In contrast, no significant differences were observed between the bare scaffolds and the Neg miR-loaded scaffolds for either the pro- or anti-inflammatory cell populations, suggesting that the observed effects were specific to miR-219/miR-338 and not attributable to RBCEVs alone.

The potential side effect of implantation-induced tissue retraction was also accessed. As shown in Figure S4, no apparent tissue retraction was observed when comparing the bare scaffolds with the RBCEVs-loaded scaffolds, as indicated by GFAP-stained astrocytes retraction distance, suggesting a favorable tissue-implant integration.

## 4. Discussion

Although gene therapy is highly promising, commonly used transfection reagents like cationic lipids and polymers often suffer from low transfection efficiency and high cytotoxicity [2]. For example, Lipofectamine-siRNA complexes are prone to forming aggregates [26], and the TransIT-TKO system loses efficacy in the presence of serum [27], both limiting their clinical utility. In contrast, RBCEVs represent a promising alternative to conventional transfection reagents due to their high delivery efficiency and low toxicity, along with immuno-evasive properties and the ability to cross biological barriers [9]. However, their applications to date have been largely confined to cancer and inflammatory disease models, with minimal exploration in the context of CNS disorders. Moreover, current strategies involving systemic administration of RBCEVs raise concerns related to off-target accumulation, poor CNS penetration, and potential systemic side effects [28]. To overcome these barriers, this work is the first to systematically assess the performance of RBCEVs on CNS neural cells in terms of cytotoxicity, uptake efficiency, and gene knockdown capability. In addition, it introduces a 3D-printed scaffold-based platform for localized and sustained sncRNA delivery via RBCEVs, offering a novel strategy for CNS-targeted RNA therapeutics.

To evaluate a potential transfection vector, the primary consideration lies in balancing cytotoxicity and transfection efficiency. In our previous work, RBCEVs demonstrated no detectable cytotoxicity when compared to commercial reagents such as Lipofectamine™ 3000 and INTERFERin®, which induced approximately 30% cell death [16]. However, their cytocompatibility in neural cells has not been assessed. When comparable amounts of knockdown-relevant dosage of RNA were used, RBCEVs resulted in similar cell viability across other neural types but preserved 10% higher viability in OPCs compared to TransIT-TKO, underscoring their superior cytocompatibility for CNS RNA delivery.

Gene knockdown efficiency is influenced by multiple factors, among which cellular uptake and intracellular trafficking of RNA-vector complexes are considered critical [14]. Our previous study systematically compared the uptake efficiency of RBCEVs across various cancer cell lines and macrophages, revealing cell type–dependent variability, with macrophages exhibiting the highest uptake capacity [14]. In the present study, primary neural cells also exhibited distinct cell type–specific uptake patterns. Microglia, which function similar to macrophages but unique in the CNS, demonstrated the highest RBCEVs uptake (∼90%), followed by OPCs at ∼76%.

Following internalization, the intracellular fate of RBCEVs namely endolysosomal escape and RNA-vector decomplexation, play pivotal roles in determining the amount of functionally active cytoplasmic RNA [29]. As shown in Figure 2D to 2F, OPCs exhibited a greater extent of both endolysosomal escape and decomplexation of RBCEVs-associated RNA compared to microglia, which explained why RBCEVs triggered higher extent of gene knockdown in OPCs despite of lower uptake than microglia, which is also consistent with other studies showing that phagocytic cells often sequester internalized cargos in lysosomes, reducing cytosolic bioavailability of functional siRNA [30, 31]. In addition, RBCEVs achieved knockdown efficiency comparable to TransIT-TKO despite a reduced miRNA load (Figure 3). This, in combination with high cytocompatibility and enhanced intracellular processing in OPCs, highlights RBCEVs as a highly suitable vector for OPCs-directed therapies.

This study also represents the first attempt to incorporate RBCEVs into a scaffold-based delivery system, with comprehensive evaluation both *in vitro* and *in vivo*. The *in vitro* evaluation of this delivery system focused on its capacity to release the cargos in a sustainable manner while preserving their bioactivity. As shown in Figure 4C, this RBCEVs-encapsulated scaffold could permit sustained release for at least 21 days, releasing 21.6% of miR at 24 h and 73.8% at Day 21. Based on the total loading amount, 48 h of incubation released ∼10 ng of siRNA, triggering ∼30% gene knockdown in OPCs (Figure 4D). In comparison, bolus transfection with 500 ng siRNA induced ∼75% gene knockdown (Figure 2C), indicating the effectiveness of scaffold-mediated delivery and the non-linear dosage-dependency of RBCEVs-mediated gene knockdown. Additionally, this sustained release over 21 days could successfully cover the myelination process of OPCs, which is evidenced in Figure B that 2 weeks co-incubation could successfully trigger enhanced OPCs differentiation and maturation. More importantly, although both the RBCEV-loaded scaffold and the previously reported one-time TKO-miR bolus transfection increased MBP⁺ cell number, myelin sheath length, and sheath count at Day 14 [21]., the scaffold-mediated delivery resulted in significantly greater enhancements across all measures, indicating a superior effect of sustained release.

The *in vivo* evaluation focused on three key aspects: (1) whether the scaffold could effectively deliver RBCEVs and RNA to OPCs, (2) whether delivery of miR-219/miR-338 could promote OPC differentiation *in vivo*, and (3) whether the scaffold would trigger undesirable immune responses.

Compared to our previous intralesional injection approach, where RBCEV signals rapidly declined within 72 hours [16], the current scaffold-based method retained a substantial amount of RBCEVs locally even after 7 days (Figure 6A). Moreover, while intravenous injection resulted in only ∼3% of EVs remaining in circulation after 12 hours [16], the scaffold-mediated delivery effectively restricted RBCEVs distribution to within 3 mm of the injury site (Figure 6A), reducing systemic exposure. These findings highlight the advantages of scaffold-based delivery in enhancing local retention and minimizing off-target distribution. Furthermore, similar to the *in vitro* uptake profile, RBCEVs were mainly taken up by microglia, OPCs, and neurons *in vivo*, with limited uptake by astrocytes, confirming a consistent preference in cellular targeting. Moreover, since miR-219 and miR-338 exert their effects at different stages of oligodendroglial-mediated myelination [25], it is essential to determine whether RBCEVs can be delivered to both undifferentiated PDGFRα⁺ OPCs and differentiated CC1⁺ oligodendrocytes to ensure comprehensive therapeutic efficacy. Current remyelination strategies primarily focus on expanding the OPCs population through protective molecule treatment like neurotrophins or stem cell transplantation [32], yet the precise regulation of OPC lineage progression remains insufficiently addressed. Despite our previous investigation on the therapeutic efficacy of miR-219/miR-338 *in vivo* [21], it was unknown whether RBCEVs-based scaffolds could deliver them to the appropriate oligodendroglial stages where their remyelination effects are most active. In this study as shown in Figure 6D and 6E, RBCEVs signals were detected in both PDGFRα⁺ OPCs and CC1⁺ mature oligodendrocytes, confirming successful RNA delivery to distinct stages of the oligodendrocyte lineage to permit the therapeutic downstream effects.

OPCs differentiation into OLs is the first step of remyelination, as characterized by the decreased expression of PDGFRα and increase expression of CC1 [33]. In this study, miR-219/miR-338 delivery led to an 85% reduction in PDGFRα⁺ cells and a two-fold increase in CC1⁺ cells, indicating effective promotion of OPC differentiation and maturation. Compared to our previous PCL/collagen-based fiber-hydrogel scaffold, which induced a moderate 50% increase in CC1⁺ cells without significantly reducing PDGFRα⁺ cells [21], the current approach exhibited a more robust differentiation-inducing effect. Given that this study applied a full-transection model, in contrast to the hemi-incision model used in the earlier work, the variation observed could potentially be attributed to injury severity, which is known to impact OPC recruitment and intrinsic regenerative capacity. Importantly, this RBCEVs-encapsulated scaffold required only 40 ng of miRs per scaffold, in contrast to the 2 μg/scaffold used in the electrospun platform. The enhanced differentiation observed despite the lower loading could be attributed to the RNA-protective capacity of RBCEVs as demonstrated in our previous studies [16], and their superior transfection efficiency in OPCs (Figure 2C). In addition, the two-fold increase in CC1⁺ cells observed *in vivo* at Day 14 is comparable to the increase in MBP⁺ cells observed *in vitro* at the same time point, indicating that the *in vitro* assay of the scaffold is highly predictive of *in vivo* performance, which holds significant relevance for future clinical applications.

The immune response is also known to significantly influence both tissue damage and regeneration processes after SCI [34]. Our previous study demonstrated that RBCEVs could modulate macrophage polarization, inducing decreased CD86 and increased CD163 expression in peripheral blood mononuclear cell (PBMCs), similar to the effects of anti-inflammatory cytokines, IL-4 and IL-10. Although CD206 was not elevated in RBCEVs-treated PBMCs, subsequent experiments showed that liver-derived F4/80⁺ macrophages exhibited increased expression of CD80, CD163, and CD206, suggesting a region- or cell type-specific immune response to RBCEVs [14] and raising the question of how CNS microglia would respond to such treatment. In the current study, scaffolds loaded with miR-219/miR-338 RBCEVs significantly increased CD206 and decreased iNOS expression in microglia, indicating a shift toward an anti-inflammatory phenotype. In contrast, scaffolds containing RBCEVs with negative control miRs elicited CD206 and iNOS levels comparable to the bare scaffold (Figure 8), confirming that the observed immunomodulation was driven by the delivered miRs rather than RBCEVs. These findings were also supported by our earlier work showing that miR-219/miR-338 suppressed pro-inflammatory cytokines, including TNF-α, IL-1 α, and IL-1β, in both *in vitro* and *in vivo* [23]. Further investigation into the underlying signaling pathways such as RNA-sequencing and cytokine-profiling is warranted to elucidate the precise mechanisms governing this miR-mediated immune modulation.

Despite all the advantages of scaffold-based delivery discussed before, integrating RBCEVs into the DLP printing process presents many challenges. First, the hemoglobin content in RBCEVs exhibits a strong absorbance peak at 400-450 nm [35], overlapping with the activation wavelength of photo-initiators such as LAP, which are commonly used in DLP printing. This leads to light absorption by RBCEVs, limiting light penetration through the bio-ink and impeding the photo-crosslinking process. Second, the nanoscale size of RBCEVs (∼200 nm) causes light scattering, adversely affecting printing resolution and structural precision [36], while also contributing to scaffold instability after implantation, as illustrated by the collapsed channels in Figure 6A. Addressing these challenges requires comprehensive investigations into the influence of RBCEVs loading on the ink printability as reflected by changes in refractive index, light absorbance, penetration depth, and X-Y plane dimension deviation from the CAD design. After establishing this, further evaluation of scaffold mechanical stiffness and biological activity would be necessary to further optimize the scaffold for potential clinical applications.

Building upon these findings, future studies should aim to further optimize RBCEVs loading and printing parameters to modulate the release profile and mechanical properties of the scaffold, thereby enhancing its regenerative capacity. In addition, long-term *in vivo* assessments are needed to evaluate the sustained impact on remyelination, which typically requires a minimum of 4 weeks post-injury [37], as well as to monitor functional recovery. Surface modification strategies could also be explored to improve the targeting efficiency of RBCEVs and enhance their delivery specificity [38]. Additionally, our previous study indicated that both miR-219/miR-338 and RBCEVs can trigger a broad range of downstream effects. To ensure clinical safety and efficacy, these effects should be thoroughly characterized in future studies as part of the translational evaluation of this delivery platform.

## 5. Conclusion

This study established a 3D-printed scaffold-based platform for the localized and sustained delivery of RBCEV-encapsulated miR-219/miR-338, with validated efficacy both *in vitro* and *in vivo*. Specifically, RBCEVs exhibited superior cytocompatibility and transfection efficiency with cell type–specific uptake, efficient endolysosomal escape, and enhanced RNA decomplexation in OPCs. The scaffold enabled sustained miRNA release over 21 days, effectively promoting OPCs differentiation and maturation *in vitro*. In a Rat complete transection model, RBCEVs were successfully delivered to both PDGFRα⁺ OPCs and CC1⁺ oligodendrocytes, supporting RNA delivery across key remyelination stages. Additionally, miR-219/miR-338 delivery modulated the immune microenvironment by shifting microglial polarization toward an anti-inflammatory phenotype. These findings highlighted the therapeutic potential of scaffold-mediated RBCEVs delivery for CNS repair and underscore the need for further optimization of printing parameters to accommodate the optical properties of RBCEVs.

## Acknowledgements

This research is supported by the Ministry of Education, Singapore, under its MOE Tier 1 (RT21/23, RG92/22) and Tier 2 grants (MOE-T2EP30220-0002). We would like to acknowledge *Neuroscience @ NTU, Interdisciplinary Graduate Programme* for providing the Nanyang Research Scholarship to carry out the research works.

