## Supplementary material for "3D-Printed Scaffolds Encapsulating Red Blood Cell Extracellular Vesicles for MicroRNA Delivery": Table S1; Figure S1; Figure S2; Figure S3; Figure S4

### Slide 1
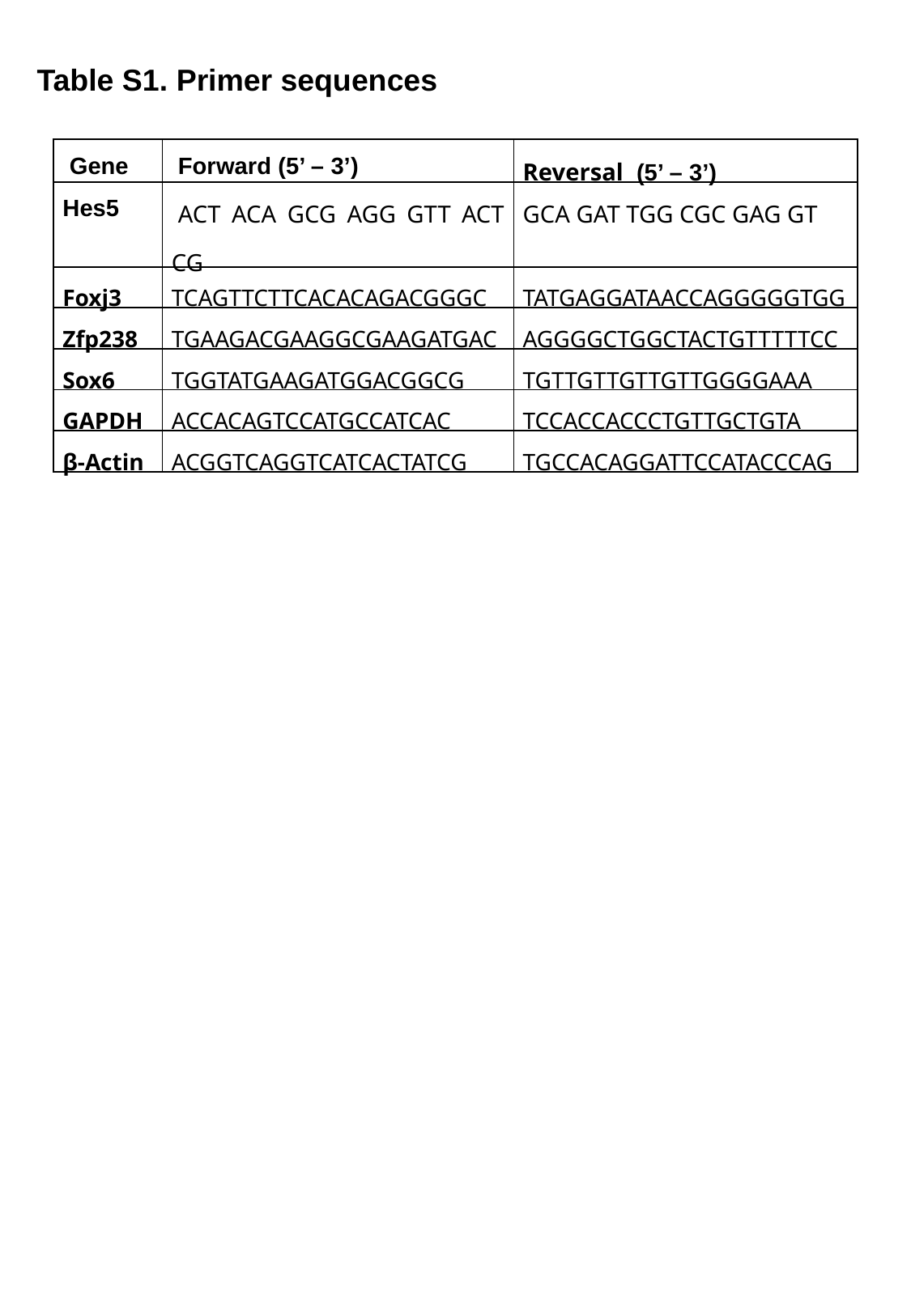

Table S1. Primer sequences
| Gene | Forward (5’ – 3’) | Reversal (5’ – 3’) |
| --- | --- | --- |
| Hes5 | ACT ACA GCG AGG GTT ACT CG | GCA GAT TGG CGC GAG GT |
| Foxj3 | TCAGTTCTTCACACAGACGGGC | TATGAGGATAACCAGGGGGTGG |
| Zfp238 | TGAAGACGAAGGCGAAGATGAC | AGGGGCTGGCTACTGTTTTTCC |
| Sox6 | TGGTATGAAGATGGACGGCG | TGTTGTTGTTGTTGGGGAAA |
| GAPDH | ACCACAGTCCATGCCATCAC | TCCACCACCCTGTTGCTGTA |
| β-Actin | ACGGTCAGGTCATCACTATCG | TGCCACAGGATTCCATACCCAG |

### Slide 2
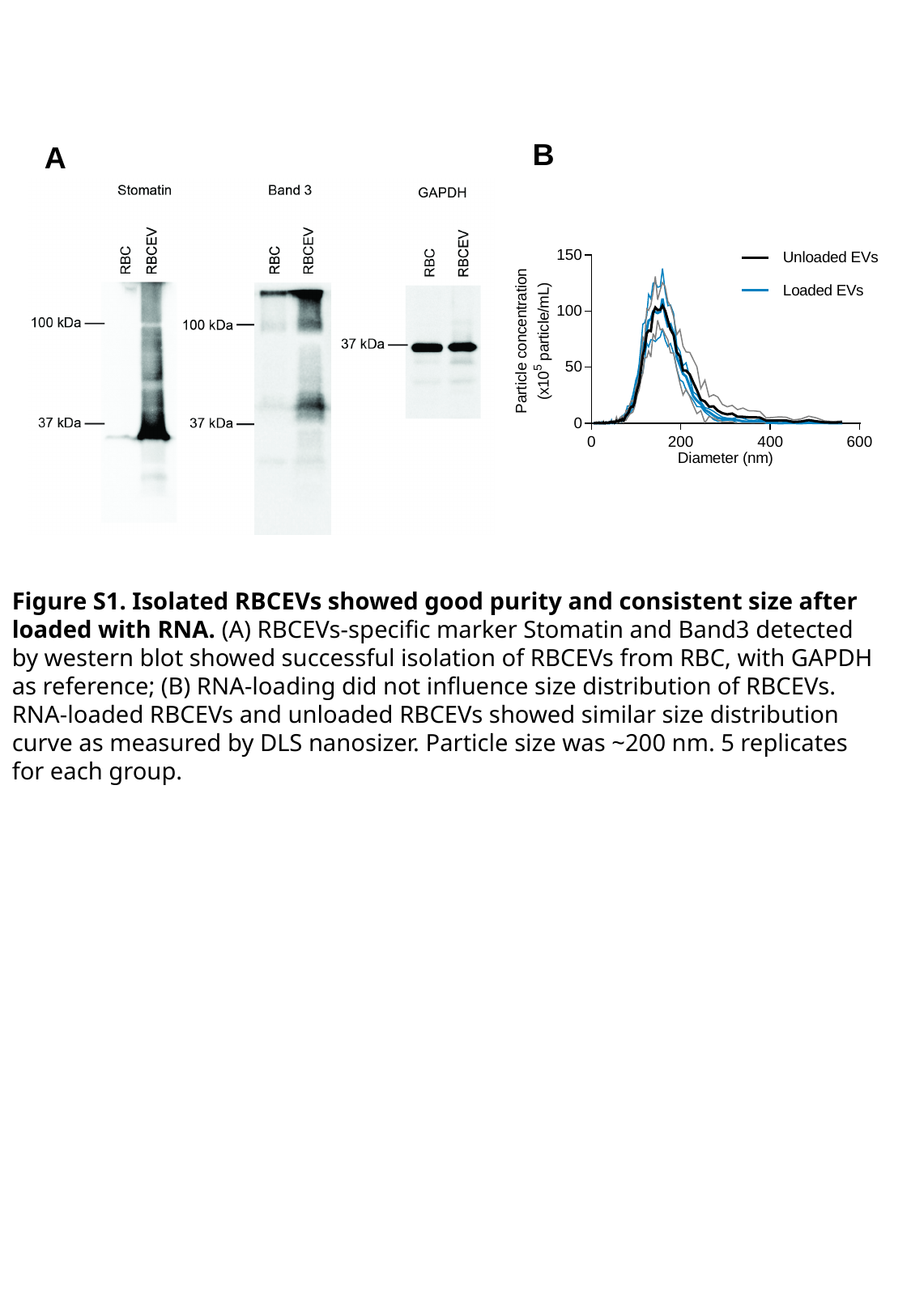

B
A
Figure S1. Isolated RBCEVs showed good purity and consistent size after loaded with RNA. (A) RBCEVs-specific marker Stomatin and Band3 detected by western blot showed successful isolation of RBCEVs from RBC, with GAPDH as reference; (B) RNA-loading did not influence size distribution of RBCEVs. RNA-loaded RBCEVs and unloaded RBCEVs showed similar size distribution curve as measured by DLS nanosizer. Particle size was ~200 nm. 5 replicates for each group.

### Slide 3
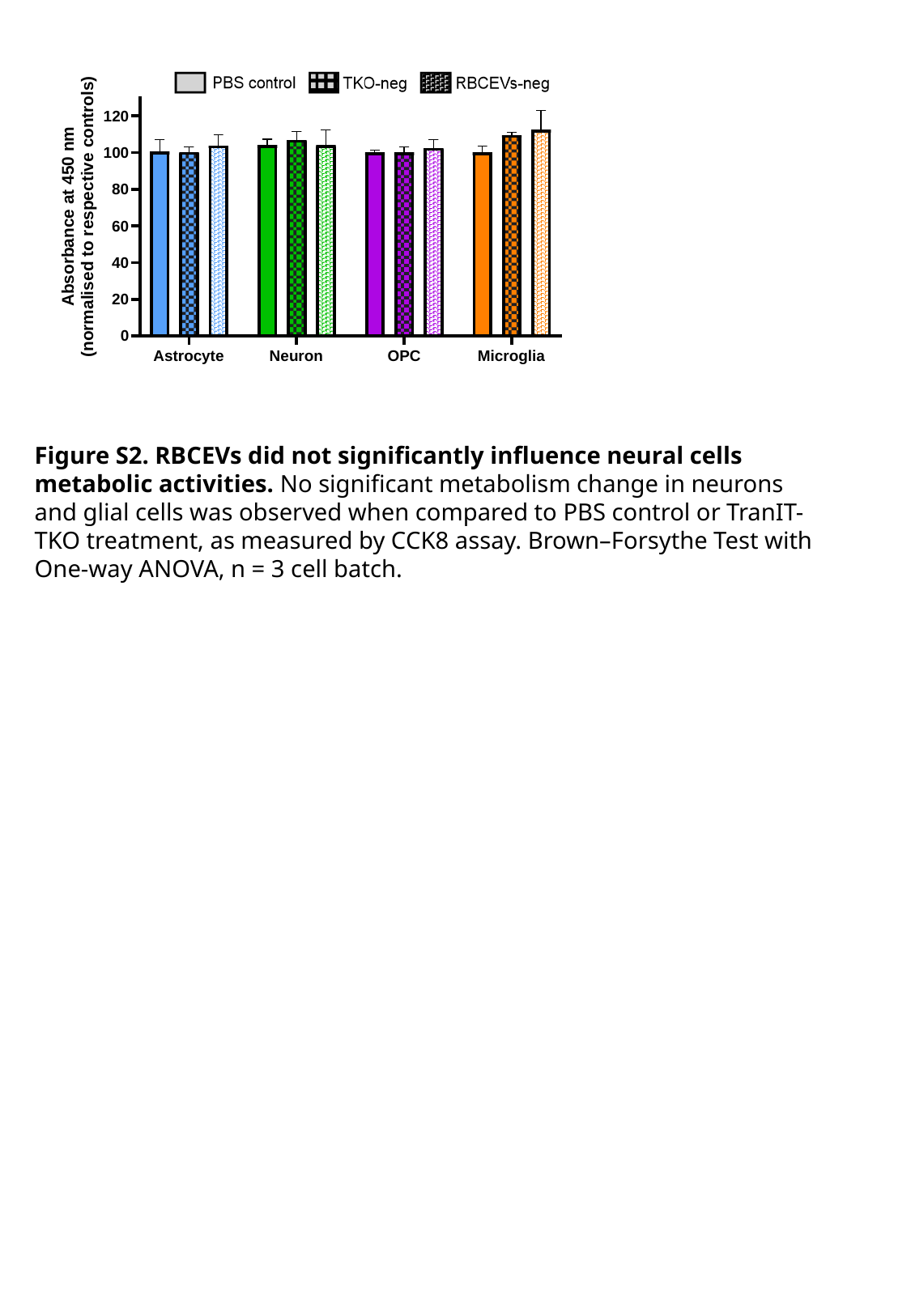

Figure S2. RBCEVs did not significantly influence neural cells metabolic activities. No significant metabolism change in neurons and glial cells was observed when compared to PBS control or TranIT-TKO treatment, as measured by CCK8 assay. Brown–Forsythe Test with One-way ANOVA, n = 3 cell batch.

### Slide 4
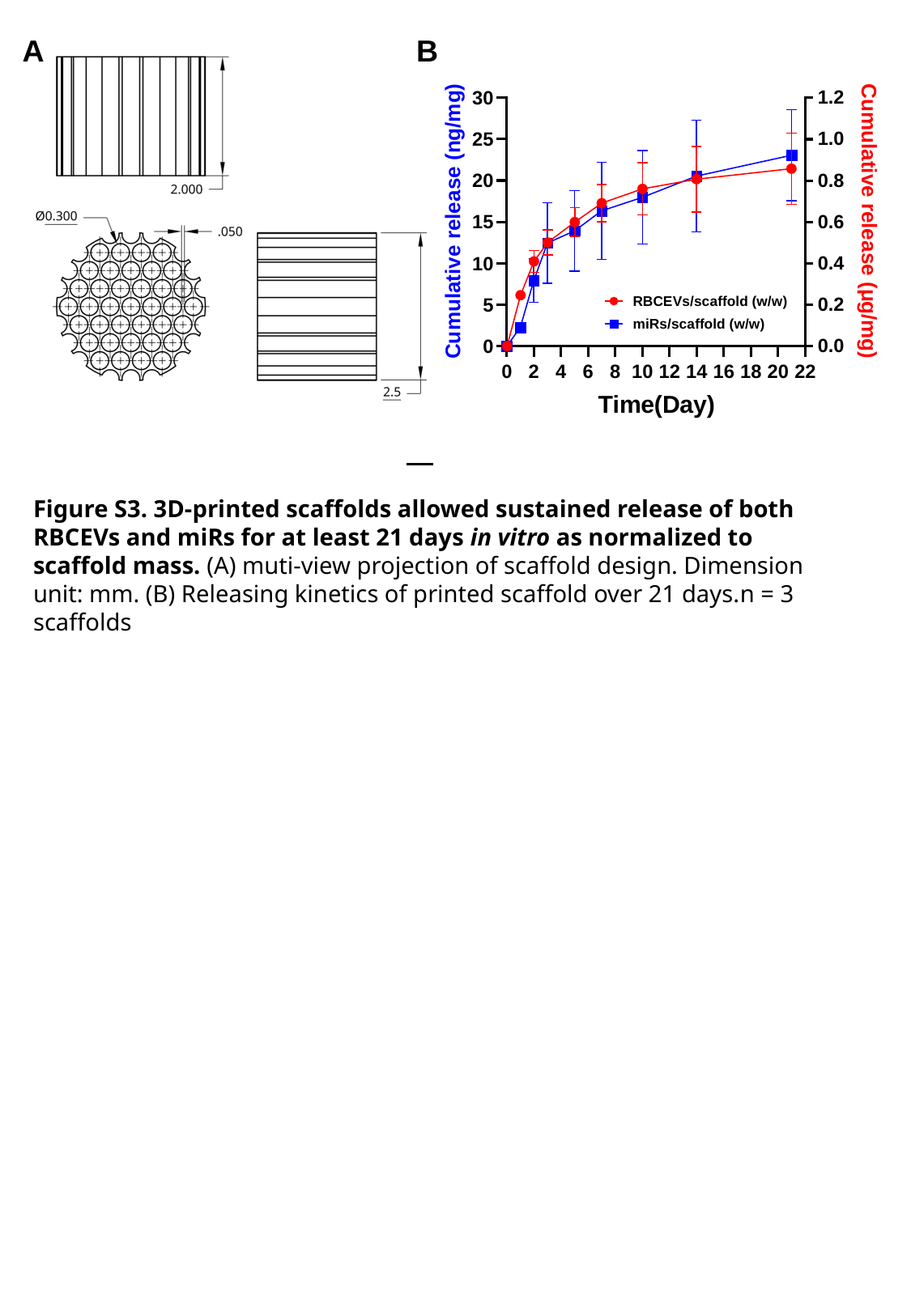

A
B
Figure S3. 3D-printed scaffolds allowed sustained release of both RBCEVs and miRs for at least 21 days in vitro as normalized to scaffold mass. (A) muti-view projection of scaffold design. Dimension unit: mm. (B) Releasing kinetics of printed scaffold over 21 days.n = 3 scaffolds

### Slide 5
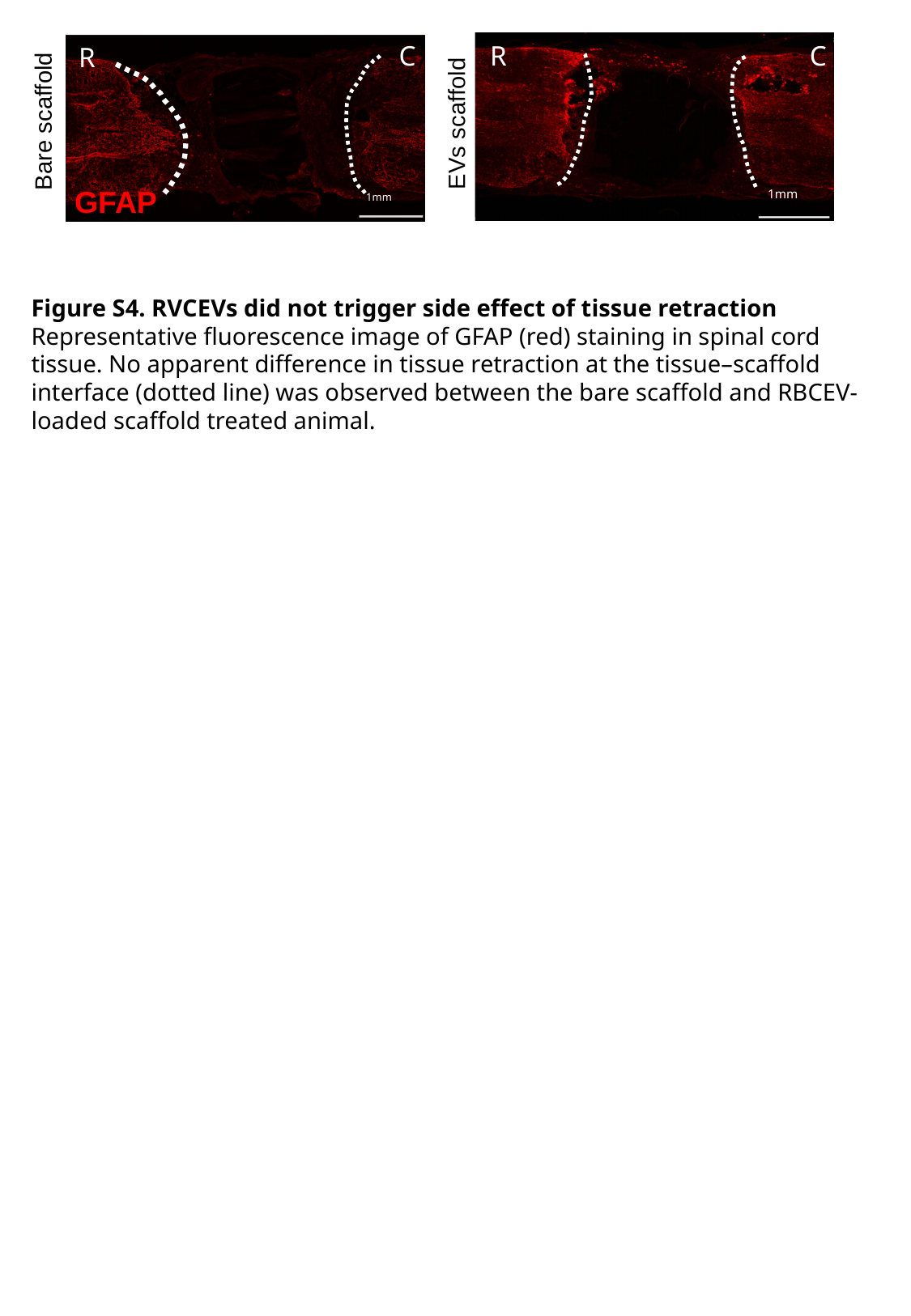

C
R
C
R
1mm
Bare scaffold
EVs scaffold
GFAP
1mm
Figure S4. RVCEVs did not trigger side effect of tissue retraction
Representative fluorescence image of GFAP (red) staining in spinal cord tissue. No apparent difference in tissue retraction at the tissue–scaffold interface (dotted line) was observed between the bare scaffold and RBCEV-loaded scaffold treated animal.
